# Cell wall remodeling is required for crowding homeostasis during cell growth in *S. cerevisiae*

**DOI:** 10.64898/2026.06.12.731398

**Authors:** Keiichiro Sakai, Marina Kunzi, Federico Uliana, Leo Willig, Ruth Lappalainen, Yoshiaki Kamada, Gabriel E Neurohr, Amy E Ikui

## Abstract

The cytoplasm is a densely crowded environment that plays essential roles in intracellular functions. Its biophysical properties can be markedly altered in response to environmental cues and changes in cellular state. Recent studies have shown that excessive growth leads to cytoplasmic dilution, contributing to cellular dysfunction. However, the molecular mechanisms regulating biophysical properties during excessive growth remain poorly understood. Here, we demonstrate that cell wall remodeling is crucial to maintain cytoplasmic crowding homeostasis during cell growth. Live-cell imaging and phosphoproteomic analysis showed that Mpk1, a component of Cell Wall Integrity (CWI) pathway, accumulates in the nucleus and its activity progressively increases as cells enlarge. Mpk1 is required to restrict cell volume expansion and maintain cytoplasmic crowding by adjusting cell wall thickness and mechanical resistance. Moreover, Mpk1 is essential for the survival of enlarged cells upon cell cycle re-entry. In addition to Mpk1, we identified multiple key components of the CWI pathway that are required to maintain physiological state of enlarged cells. Together, these findings establish the CWI signaling pathway as a key regulator of cell size homeostasis and cytoplasmic biophysical properties, providing a mechanistic link between cell wall remodeling and intracellular properties.

## Introduction

In normally proliferating cells, the cytoplasm is a highly crowded environment densely packed with macromolecules ^1^. Macromolecular crowding can promote intermolecular assembly by increasing effective concentrations and favoring compact associated states, although it may also impede molecular diffusion ^1–3^. Accumulating evidence indicates that cytoplasmic biophysical properties influence a wide range of intracellular processes, including kinase-mediated signaling ^4–7^, phase separation ^8,9^, protein production ^10^, and microtubule dynamics ^11^. Furthermore, cytoplasmic biophysical properties are dynamically altered in response to various environmental conditions and cellular states ^8,12–16^. These alterations can potentially cause catastrophic failure of cellular system by disrupting essential biological processes. To preserve cytoplasmic homeostasis, several regulatory mechanisms have been identified; one of the well-characterized examples is WNK1, a molecular crowding sensor that detects and adjusts cytoplasmic properties under hyperosmotic stress ^17^.

Recent evidence suggests that excessive cell growth leads to cytoplasmic dilution, resulting in a less crowded cytoplasmic environment ^18–21^. Cells normally maintain their size within a defined range, and deviations from this range are often associated with cellular dysfunction ^22–24^. The consequences of cell enlargement have been studied in both yeast and mammalian systems, where enlarged cells can be generated by uncoupling cell growth from cell division. Cytoplasmic dilution in enlarged cells is thought to arise from reduced concentrations of macromolecules, including RNA, proteins, and ribosomes, which normally serve as major cytoplasmic crowders ^8,9,18,25,26^. Such dilution may result from a declining DNA-to-cytoplasm ratio as cells enlarge, causing DNA-dependent biosynthetic capacity to become limiting for macromolecule production ^18,21,27^.

Reduced cytoplasmic crowding in enlarged cells is likely to broadly impair intracellular organization and biochemical function. Indeed, enlarged yeast cells are accompanied by global changes in protein concentrations, selective scaling defects across the proteome, and dilution of key cellular components, demonstrating that increased size directly alters molecular composition and physiological state ^18,19,28–31^. Parallel observations have emerged in mammalian cells. For example, chemical perturbation of cell cycle progression allows cells to continue growth without division, producing enlarged cells with hallmarks of senescence ^32^. Enlarged mammalian cells also display proteomic remodeling, impaired signaling, and senescence-associated phenotypes ^28,33–37^. Together, these studies support a conserved principle that growth beyond the normal cell-size range disrupts molecular homeostasis and progressively compromises cellular function in eukaryotes. However, the mechanisms that sense and respond to changes in cell size and cytoplasmic biophysical properties remain poorly understood.

In walled cells such as yeasts, cell volume expansion during growth is driven by high internal hydrostatic pressure, known as turgor pressure, which pushes the plasma membrane against the cell wall ^38–41^. This pressure generates substantial mechanical stress and can produce sufficient membrane tension to cause cell lysis if not properly constrained. The cell wall therefore serves a critical role in counteracting turgor pressure by providing mechanical reinforcement and preserving cellular integrity. When the balance between turgor pressure and the mechanical properties of the cell wall is disrupted, the rate of volume expansion may become abnormally accelerated or impaired. Notably, in spherical cells the turgor-derived stress on the cell wall is predicted to increase with cell size by Laplace’s law, a model previously applied by others ^19,40,42,43^. However, it remains unclear how cells cope with this increased stress to maintain homeostasis when cell size increases during growth.

In this study, we demonstrate that the Cell Wall Integrity (CWI) pathway maintains both cell size and cytoplasmic properties in the budding yeast *Saccharomyces cerevisiae* during cell growth. Phosphoproteomic analysis revealed that Mpk1, a mitogen-activated protein kinase (MAPK) in the CWI pathway, becomes progressively activated as cell size increases. Loss of key CWI components in enlarged cells resulted in further cell enlargement, reduced cytoplasmic crowding, and failure in cell cycle re-entry, leading to cell lysis. Thus, the CWI pathway restrains excessive cell growth while maintaining cytoplasmic concentration. Mechanistically, Mpk1 promotes cell wall stiffness and mechanical resistance in enlarged cells, consistent with predictions from Laplace’s law. This mechanism provides a novel avenue for regulating cell size and modulating cytoplasmic crowding in a cell size-dependent manner.

## Results

### The mobility of 40 nm particles in the cytoplasm increases with cell size

To investigate the relationship between cell size and cytoplasmic crowding in budding yeast, we employed cell cycle mutants that exhibit smaller (*whi5*Δ) or larger (*cln3*Δ) cell size compared with wild-type cells (Fig. 1A) ^28,44^. The mean cell volumes of the *whi5*Δ, wild-type, and *cln3*Δ strains were 47, 54, and 95 fL, respectively (Fig. 1B). To evaluate cytoplasmic crowding in these cells, we used genetically encoded multimeric nanoparticles (GEMs) that self-assemble into 40 nm spherical particles, hereafter referred to as 40nm-GEMs ^8^. The gene encoding 40nm-GEMs fused to a fluorescent protein T-Sapphire was integrated at a defined genomic locus and resulted in the stable assembly of bright tracer particles in the cytoplasm (Fig. 1A). We tracked the 40nm-GEMs in a live cell to determine the intracellular trajectories and computed the effective diffusion coefficient (D_eff_) using mean-square displacement (MSD) analysis. To assess the 40nm-GEMs particle mobility relative to cell size across strains, the median D_eff_ was plotted as a function of mean cell volume measured by a Coulter counter (Fig. 1C). The median D_eff_ of *cln3*Δ cells was higher than that of *whi5*Δ and wild-type cells, consistent with their increased cell size (Fig. 1C). We also computed the median D_eff_ at single cell level and plotted as a function of cell area measured based on the bright-field images (Fig. 1D). There was a clear correlation between particle mobility and cell size in *whi5*Δ, wild-type, and *cln3*Δ cells (Fig. 1D). These results indicate that cytoplasmic crowding is reflected by cell size in budding yeast, in agreement with previous findings in fission yeast and mammalian cells ^18,21^.

**Fig. 1.**
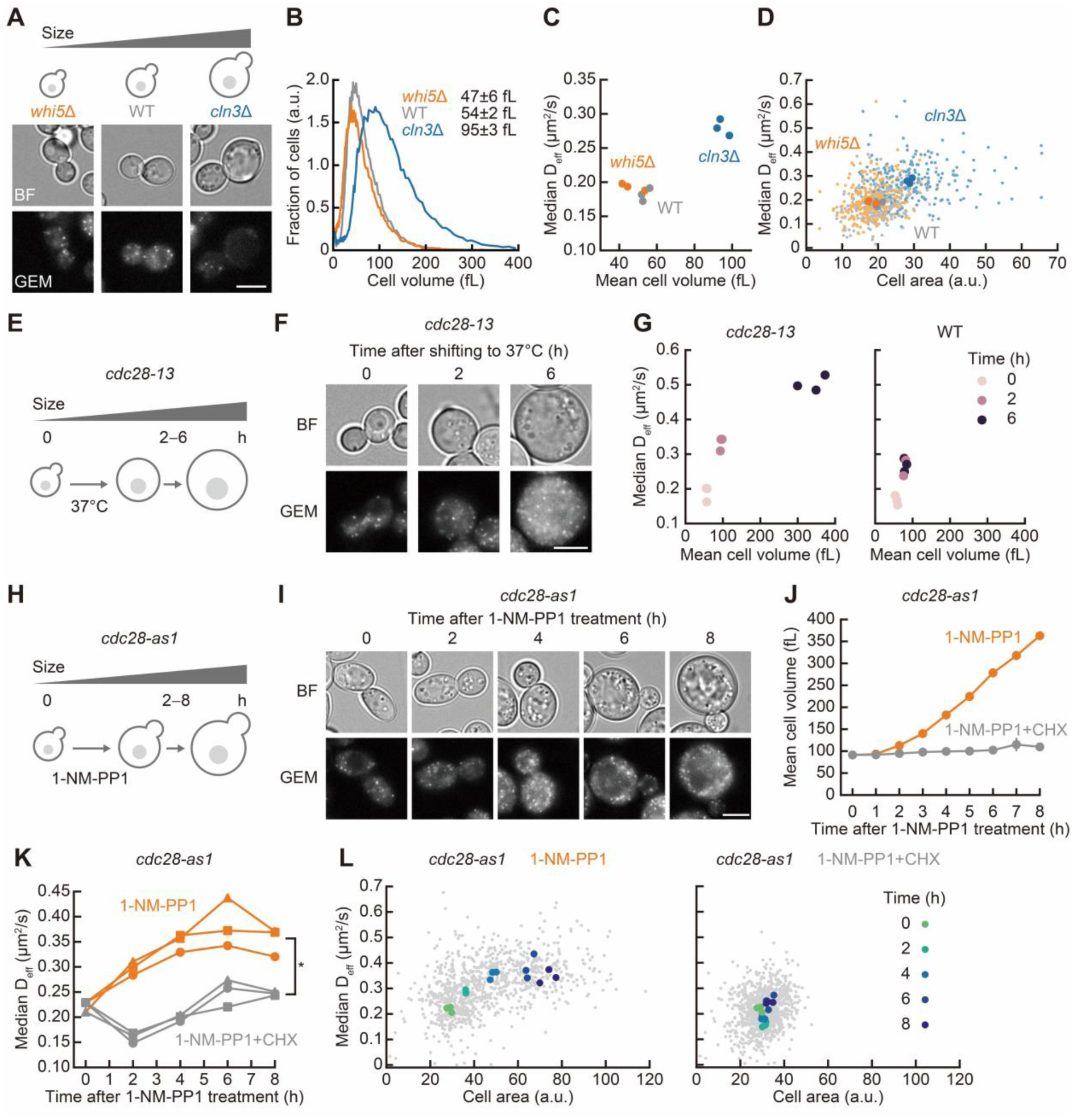
The mobility of 40 nm particles increases with cell size in budding yeast. (A) Representative bright-field (BF) and fluorescence images of *whi5*Δ, wild-type (WT), and *cln3*Δ cells expressing 40nm-GEMs. Scale bar, 5 μm. (B) Distribution of cell volume determined by a Coulter counter in *whi5*Δ (orange), WT (gray), and *cln3*Δ (blue) cells. The mean values from the three independent experiments were represented with SD. (C) Median D_eff_ of 40nm-GEMs versus mean cell volume in *whi5*Δ, WT, and *cln3*Δ cells from three independent experiments (n > 1350 trajectories for each experiment). Mean cell volumes were determined using a Coulter counter. (D) Median D_eff_ of 40nm-GEMs and cell areas at the single-cell level in *whi5*Δ, WT, and *cln3*Δ cells. Each small dot represents the median D_eff_ of 40nm-GEMs in a single cell and the cell area measured in the same cell (n > 200 cells from three independent experiments). Each large dot indicates the median D_eff_ of 40nm-GEMs and the median cell area from each experiment. Cell areas were measured from bright-field images. (E) A schematic illustration of *cdc28-13* cells after shifting to the restrictive temperature 37℃. (F) Representative BF and fluorescence images of *cdc28-13* cells expressing 40nm-GEMs after shifting to 37℃. Scale bar, 5 μm. (G) Median D_eff_ of 40nm-GEMs versus mean cell volume after shifting to 37℃ in WT and *cdc28-13* cells from three independent experiments (n > 2000 trajectories at each time point for each experiment). Mean cell volumes were determined using a Coulter counter. (H) A schematic illustration of *cdc28-as1* cells after 1-NM-PP1 treatment. (I) Representative BF and fluorescence images of *cdc28-as1* cells expressing 40nm-GEMs after 1-NM-PP1 treatment at 10 μM. Scale bar, 5 μm. (J) Cell volume determined by a Coulter counter in *cdc28-as1* cells after 1-NM-PP1 treatment. Cells were incubated with 1-NM-PP1 alone (orange) or 1-NM-PP1 with cycloheximide (1-NM-PP1+CHX, gray). The mean values from the three independent experiments were represented with SD. (K) Median D_eff_ of 40nm-GEMs in *cdc28-as1* cells treated with 1-NM-PP1. The values from three independent experiments are indicated by circles, squares, and triangles, respectively (n > 1870 trajectories at each time point for each experiment). Statistical significance was assessed using a two-tailed Welch’s t-test at 8 h. *, p < 0.05. (L) Median D_eff_ of 40nm-GEMs and cell areas at the single-cell level for *cdc28-as1* cells treated with 1-NM-PP1. Each small gray dot represents the median D_eff_ of 40nm-GEMs in a single cell and the cell area measured in the same cell (n > 100 cells at each time point from three independent experiments). Each large dot indicates the median D_eff_ of 40nm-GEMs and median cell area at each time point obtained from each experiment. Cell areas were measured from bright-field images.

To induce excessive cell growth in a conditional manner, we used a temperature-sensitive allele of *CDC28*, the yeast homolog of human *CDK1* (*cdc28-13* mutant). The *cdc28-13* cells arrest the cell cycle in G1 upon shifting the temperature from 26℃ to 37℃, enabling precise control over cell size with a wide size distribution (Fig. 1E) ^18,31,45^. The *cdc28-13* cells expressing 40nm-GEMs were incubated at 37℃ for up to 6 h, resulting in an increase in cell size with bright tracer particles (Fig. 1F). Flow cytometry analysis confirmed an increase in G1-peak at 2 and 6 h after the shift to 37℃ in *cdc28-13* cells, which was not observed in wild-type cells (Fig. S1A). The particle mobility progressively increased in *cdc28-13* cells over 6 h, which correlated with cell volume expansion, whereas there were no substantial changes in either particle mobility or cell volume in wild-type cells (Fig. 1G). These results suggest that cytoplasmic crowding is decreased in enlarged *cdc28-13* cells, as reported previously ^19^.

We next employed an analog-sensitive mutant *cdc28-as1*, which arrests the cell cycle either at G1 (unbudded cells) or G2/M (small budded cells) when treated with the nonhydrolyzable ATP analog 1-NM-PP1 (Fig. 1H and S1B) ^46^. The budding index only mildly changes after inhibitor addition, showing that *cdc28-as1* cells arrest the cell cycle at various stages (Fig. S1C). In the absence of 1-NM-PP1, *cdc28-as1* cells maintained their cell size comparable to wild-type cells (Fig. S1D). After 1-NM-PP1 treatment, *cdc28-as1* cells expressing 40nm-GEMs gradually increased and reached approximately a 3.5-fold cell volume expansion after 8 h (Fig. 1I and J, Movie S1). The particle mobility increased over 6 h, with no further increase thereafter (Fig. 1K). To determine whether the accelerated particle mobility in *cdc28-as1* cells resulted from cell enlargement rather than prolonged Cdc28 inhibition, cells were treated with the translation inhibitor, cycloheximide (CHX), to block cell growth. The cell volume expansion in *cdc28-as1* was completely inhibited by CHX in the presence of 1-NM-PP1 treatment (Fig. 1J), and the increase in particle mobility was dramatically suppressed throughout the 8 h time course (Fig. 1K). At the single cell level, 1-NM-PP1 treatment alone led to an increase in the median D_eff_ with cell size up to 4–6 h, whereas no substantial changes in either particle mobility or cell size were observed with combined 1-NM-PP1 and CHX treatment (Fig. 1L). Together, these measurements show that cell enlargement reduces macromolecular crowding, altering the biophysical properties of the cytoplasm.

### Mpk1-dependent phosphorylation progressively increases during cell enlargement

Cytoplasmic crowding impacts essential cellular processes, including protein production and cytoskeletal dynamics, and is typically maintained within a narrow range ^1,10,11,47^. We therefore asked whether cells actively regulate cytoplasmic crowding during cell growth. To identify signaling pathway that might contribute to crowding homeostasis, we performed phosphoproteomic profiling using *cdc28-13* cells at three different time points (2, 4, and 6 h) after the shift to 37℃ to monitor temporal changes in phosphorylation sites. After cell lysis and protein digestion, the resulting peptides were either enriched for phosphopeptides or analyzed directly to normalize phosphorylation levels to total protein abundance ^48^. This approach enabled the quantification of 4,985 phosphosites, of which 1,548 changed significantly in at least one condition (Fig. S2A–E). The phosphorylation and protein levels are deposited on PRIDE (PXD075955).

To identify signaling pathways that are affected by cell size, we performed clustering analysis of phosphosite profiles, considering only sites that change in at least one condition and using values normalized to total protein abundance. This analysis identified eight distinct clusters that maximize separation between groups (Fig. 2A and S2F). We subsequently performed kinase–substrate enrichment analysis (KSEA) for each cluster to predict the kinases responsible for the observed phosphorylation patterns using the NetworKIN algorithm (Fig. 2B) ^49^. Among the clusters, cluster 4 attracted our attention because it is characterized by a gradual increase in phosphorylation over time. Several phosphosites within this cluster are significantly linked to the kinase Mpk1 (p-value = 0.003, hypergeometric test) (Fig. S2G). Additional evidence supports Mpk1 activation over the time course. In particular, dual phosphorylation of threonine and tyrosine residues (T190 and Y192) within the activation loop, increases over time, indicating progressive activation of Mpk1 (Fig. 2C) ^50,51^.

**Fig. 2.**
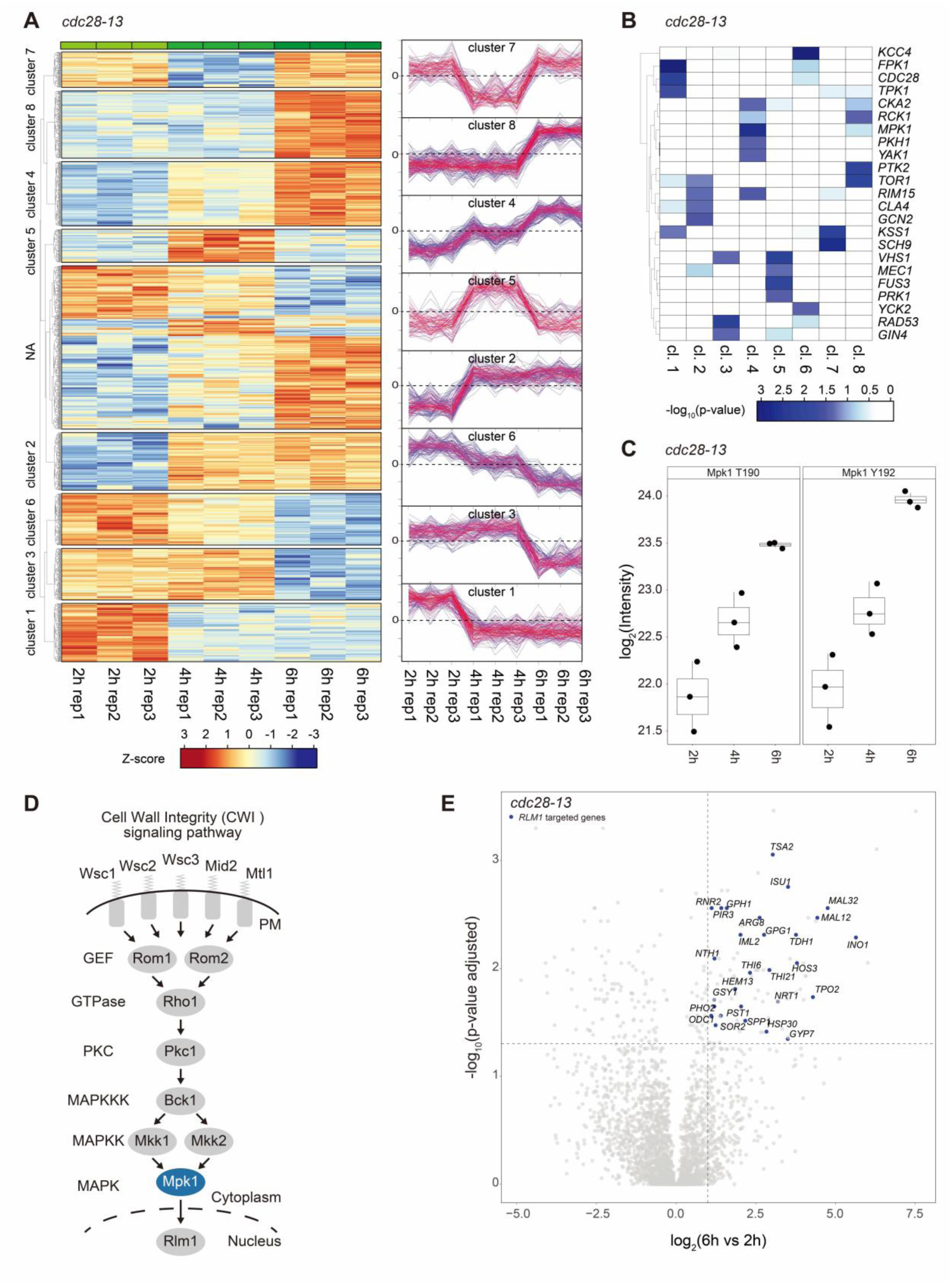
Phosphoproteomic analysis shows a progressive increase in Mpk1 kinase activity. (A) Heatmap showing phosphosite intensities normalized to protein abundance. For the 1,548 significantly changing phosphosites, values were Z-score normalized across samples. The color scale indicates relative phosphorylation levels, with red representing increased phosphorylation and blue decreased phosphorylation relative to the mean. Unsupervised hierarchical clustering identified eight clusters (cluster membership ≥ 0.6). The average Z-score profiles for each cluster are shown next to the heatmap. (B) Kinase activity was inferred for each cluster using the NetworKIN algorithm ^49^. The top 10 predicted kinases per cluster with enrichment significance (p-value < 0.05) in at least one cluster are shown. The blue color scale indicates the statistical significance of the enrichment for each kinase–cluster association. (C) Phosphorylation levels of Mpk1 at T190 and Y192 within the activation loop across the analyzed time points. (D) A schematic illustration of the cell wall integrity (CWI) pathway in budding yeast (PM; plasma membrane, GEF; guanosine nucleotide exchange factor, PKC; protein kinase C, MAPKKK; MAP kinase kinase kinase, MAPKK; MAP kinase kinase, MAPK; MAP kinase). (E) Volcano plot showing differential protein abundance between 6 h and 2 h. The x-axis represents the log₂ fold change, while the y-axis shows the p-value adjusted with Benjamini-Hochberg correction. Proteins annotated as Rlm1 target genes (log_2_ fold change > 1 and p-value < 0.05) are highlighted in blue (p-value = 1.26×10⁻⁹, hypergeometric test) ^60^.

Mpk1 is a key MAPK in the CWI pathway (Fig. 2D) ^52,53^. This pathway is activated in response to various cell wall stresses, such as detergent treatment and compressive mechanical stress, to promote cell wall repair and reinforcement ^54–56^. These stresses are sensed by mechanosensors on the plasma membrane, including Wsc1, Wsc2, Wsc3, Mid2, and Mtl1 ^52,53^. These sensors activate the MAPK signaling cascade Bck1-Mkk1/Mkk2-Mpk1 via GTPase Rho1 and protein kinase C Pkc1 ^52,53^. The Mpk1-activated transcription factor Rlm1 then regulates gene expression involved in cell wall biogenesis, contributing to the maintenance of wall integrity. Given that Mpk1 directly activates Rlm1 ^57–59^, we therefore used the global proteome dataset to examine whether Rlm1 target proteins are upregulated during cell enlargement. Among the proteins significantly more abundant at 6 h compared with 2 h, we identified several known Rlm1 targets (Fig. 2E) ^60^. Taken together, the integration of phosphoproteomic and proteomic data show a progressive activation of CWI signaling pathway during cell enlargement.

### Cell enlargement promotes nuclear localization of Mpk1

Previous studies have demonstrated that Mpk1 increases its expression levels and undergoes protein relocalization upon heat shock and mechanical compressive stresses ^54,57,58,61,62^. These Mpk1 localization changes are considered reliable indicators of CWI signaling pathway activation^52^. Our proteomic analysis revealed that Mpk1 abundance gradually increases over time in *cdc28-13* cells (Fig. S2H). To further evaluate this at a single cell level, we examined Mpk1 protein abundance and localization during cell enlargement under fluorescence microscopy. Mpk1 fused to mNeonGreen (Mpk1-mNG) was localized to both the nucleus and the cytoplasm in *cdc28-as1* cells, without 1-NM-PP1 treatment, as expected (Fig. 3A) ^54,61,62^. Upon 1-NM-PP1 treatment, the fluorescence intensity of Mpk1-mNG increased over time in both the nucleus and the cytoplasm as cell size increased (Fig. 3A). To exclude the possibility that the cellular environment, such as macromolecular crowding and viscosity, affects the fluorescence intensity of fluorescent proteins ^63^, we examined the protein expression of histone H2B (Htb1) fused to mNeonGreen (Htb1-mNG) as a relative control. The Htb1-mNG concentration gradually decreased over time upon 1-NM-PP1 treatment (Fig. S3A), consistent with previous proteome analyses showing that histone proteins abundance do not scale with increasing size during cell growth ^28^. Quantitative analysis of nuclear median fluorescence intensities revealed that Mpk1-mNG increased at ∼1.8–2.2-fold, whereas Htb1-mNG decreased at ∼0.75-fold at 8 h after 1-NM-PP1 treatment (Fig. S3B). Collectively, these data indicate that the total nuclear Mpk1 concentration increases progressively during cell enlargement, suggesting CWI signaling pathway activation in enlarged cells.

**Fig. 3.**
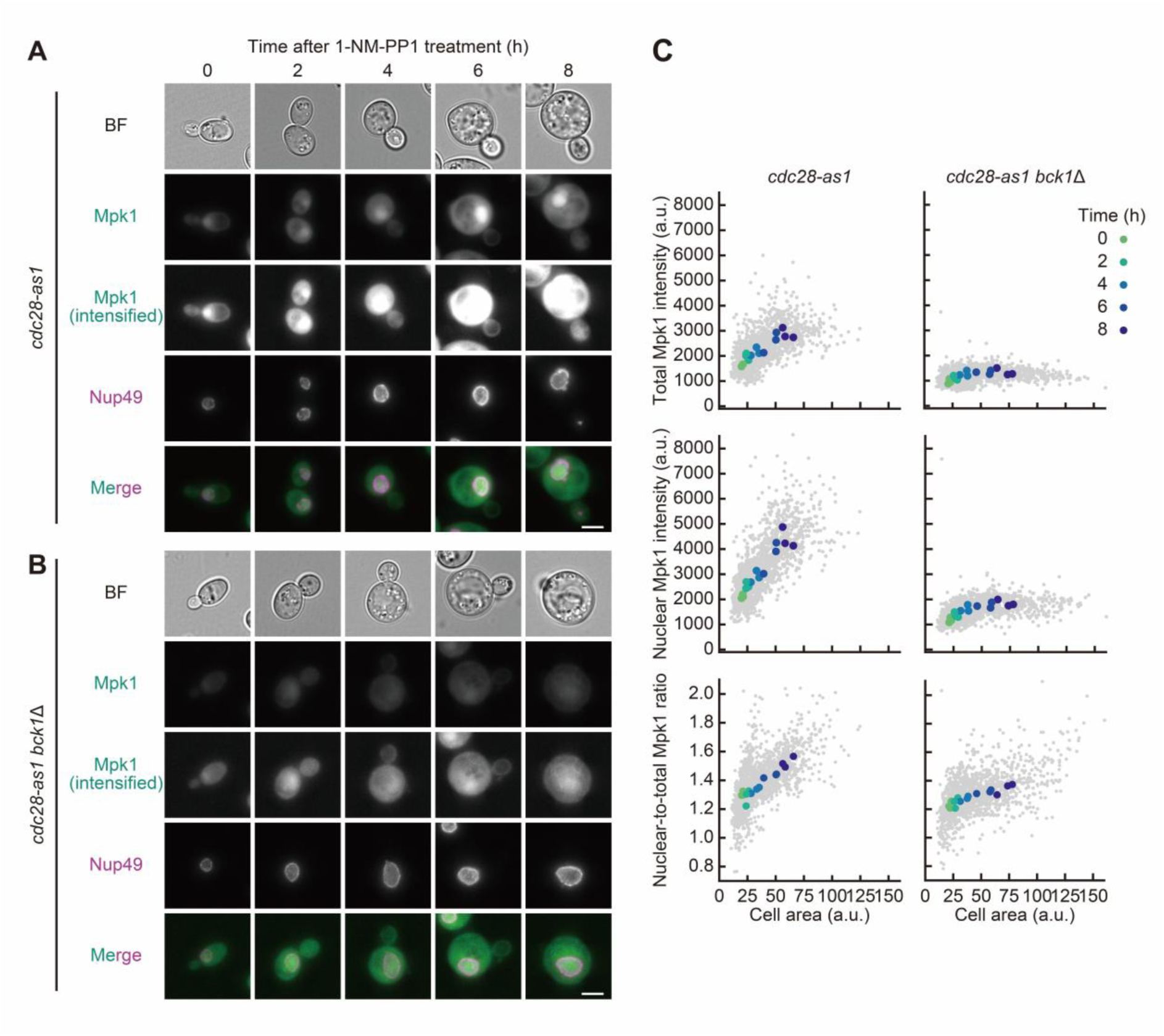
Mpk1 increases in abundance and accumulates in the nucleus during cell enlargement. (A–B) Representative images of Mpk1-mNeonGreen in *cdc28-as1* (A) or *cdc28-as1 bck1*Δ (B) cells after 1-NM-PP1 treatment for the indicated times, shown with bright-field (BF) and Nup49-mScarlet-I (nuclear marker) images. Merged images of Mpk1-mNeonGreen with Nup49-mScarlet-I are also shown. Because the Mpk1-mNeonGreen fluorescence in *cdc28-as1 bck1*Δ cells was dimmer compared to that in *cdc28-as1* cells, the intensified images are shown. For fluorescent protein signals, maximal intensity projection images are presented. Scale bar, 5 μm. (C) Mpk1-mNeonGreen intensities in whole cells (upper) or nuclei (middle) plotted against cell area at the single-cell level for *cdc28-as1* (left) and *cdc28-as1 bck1*Δ (right) treated with 1-NM-PP1. Nuclear-to-total intensity ratios are also shown (lower). Each small gray dot represents the mean Mpk1 intensity or ratio with the cell area measured in the same cell (n > 250 cells at each time point from three independent experiments). Each large-colored dot indicates the median Mpk1 intensity or ratio at each time point obtained from each experiment. Cell areas were measured from fluorescence images of Mpk1-mNeonGreen.

Mpk1 kinase activation is mediated by phosphorylation at T190 and Y192, which is dependent on the upstream MAPKKK Bck1 (Fig. 2D) ^62,64^. Interestingly, overall Mpk1-mNG intensity in *cdc28-as1 bck1*Δ cells was substantially lower than that in *cdc28-as1* cells throughout the time course (Fig. 3B). We next quantified total and nuclear Mpk1-mNG intensity as a function of cell area in *cdc28-as1* and *cdc28-as1 bck1*Δ cells (Fig. 3C). The nuclear pore protein Nup49-mScarlet-I was used as a nuclear marker. Both total and nuclear Mpk1-mNG intensities increased as cell size expanded in *cdc28-as1* cells, while deletion of *BCK1* substantially reduced both total and nuclear signals close to the basal levels (Fig. 3C). The nuclear-to-total Mpk1 intensity ratio increased as cell size becomes large in *cdc28-as1* cells, indicating the nuclear accumulation of Mpk1 (Fig. 3C). Although the nuclear-to-total ratio slightly increased with cell size in *cdc28-as1 bck1*Δ cells, it remained lower than that in *cdc28-as1* cells (Fig. 3C). These results suggest that Bck1 promotes Mpk1 nuclear accumulation during cell enlargement and together they demonstrate a progressive activation of the CWI pathway during cellular enlargement.

### Mpk1 is required to prevent excessive cell enlargement and increased particle mobility

To test whether Mpk1 is involved in cell size regulation during continuous cell growth, we measured cell size in *cdc28-as1* cells with or without *MPK1* gene. While the cell volume in *cdc28-as1 mpk1*Δ cells was comparable to that in *cdc28-as1* cells in the absence of 1-NM-PP1 (Fig. S4A), deletion of *MPK1* accelerated cell volume increase upon inhibitor addition (Fig. 4A and B). CHX treatment prevented growth of *cdc28-as1 mpk1*Δ cells in the presence of 1-NM-PP1 (Fig. 4B). We next monitored the mobility of 40nm-GEMs, which was markedly increased in *cdc28-as1 mpk1*Δ cells (Movie S2), showing significantly higher median D_eff_ at 8 h than that in *cdc28-as1* cells (Fig. 4C). The enhanced particle mobility in *cdc28-as1 mpk1*Δ cells was efficiently suppressed in the presence of CHX (Fig. 4C). Single-cell analysis also revealed a cell size-dependent increase in particle mobility in *cdc28-as1 mpk1*Δ cells compared with *cdc28-as1* cells, which was completely suppressed under CHX-treated condition (Fig. 4D). There were no obvious differences in cell volume and particle mobility between wild-type and *mpk1*Δ cells (Fig. S4B–D). We conclude that Mpk1 is required to maintain cell size and cytoplasmic crowding during excessive growth.

**Fig. 4.**
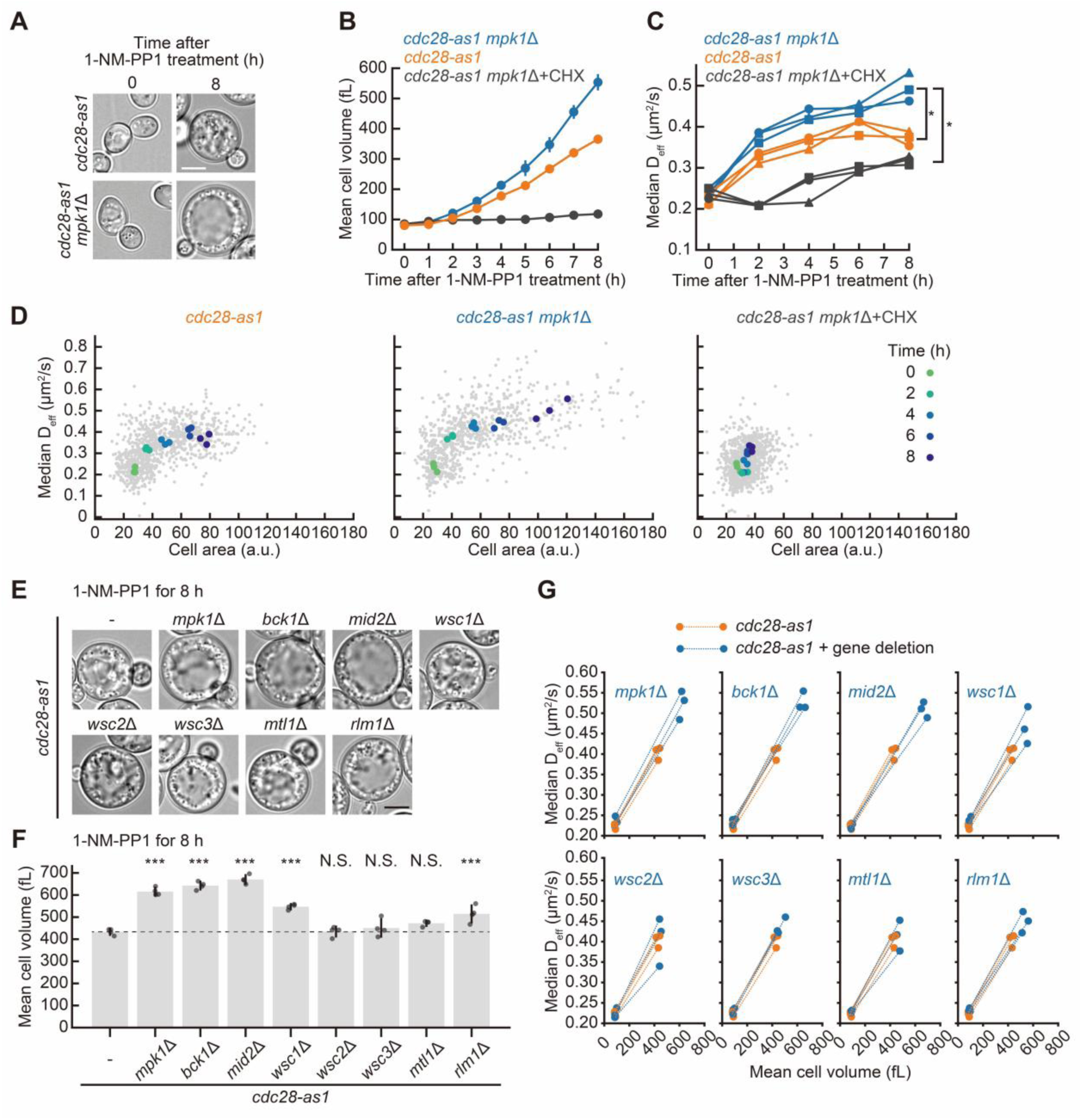
The signaling molecules in the CWI pathway maintain cell size and particle mobility during excessive growth. (A) Representative bright-field images of *cdc28-as1* and *cdc28-as1 mpk1*Δ cells after 1-NM-PP1 treatment for 8 h. Scale bar, 5 μm. (B) Cell volume in *cdc28-as1* and *cdc28-as1 mpk1*Δ cells after 1-NM-PP1 treatment. *cdc28-as1 mpk1*Δ cells were incubated with 1-NM-PP1 alone (blue) or 1-NM-PP1 with cycloheximide (1-NM-PP1+CHX, dark gray). *cdc28-as1* cells were incubated with 1-NM-PP1 alone (orange). Cell volume was measured by a Coulter counter. The mean values from the three independent experiments are represented with SD. (C) Median D_eff_ of 40nm-GEMs in *cdc28-as1* and *cdc28-as1 mpk1*Δ cells treated with 1-NM-PP1. The values from three independent experiments are indicated by circles, squares, and triangles, respectively (n > 1600 trajectories at each time point for each experiment). Statistical significance was assessed using a two-tailed Welch’s t-test with Bonferroni correction at 8 h. *, p < 0.05. (D) Median D_eff_ of 40nm-GEMs and cell areas at the single-cell level for *cdc28-as1* and *cdc28-as1 mpk1*Δ cells treated with 1-NM-PP1. Each small gray dot represents the median D_eff_ of 40nm-GEMs in a single cell and the cell area measured in the same cell (n > 100 cells at each time point from three independent experiments). Each large dot indicates the median D_eff_ of 40nm-GEMs and median cell area at each time point obtained from each experiment. Cell areas were measured from bright-field images. (E) Representative bright-field images of indicated strains after 1-NM-PP1 treatment for 8 h. Scale bar, 5 μm. (F) Cell volume of indicated strains after 1-NM-PP1 treatment for 8 h. Cell volume was measured by a Coulter counter. The mean values from the four independent experiments were represented with SD. Each dot represents the value from an individual experiment. Statistical significance was assessed by comparison with *cdc28-as1* (-) cells using one-way ANOVA followed by a *post hoc* Dunnett’s test. ***, p < 0.001; N.S., not significant. (G) Median D_eff_ of 40nm-GEMs with mean cell volume before and after 1-NM-PP1 treatment for 8 h. The values from three independent experiments are indicated (n > 2120 trajectories at each time point for each experiment). Mean cell volumes were determined using a Coulter counter.

We also employed the LexOpr-*CLN1 cln1*Δ *cln2*Δ *cln3*Δ strain, which conditionally expresses a single G1 cyclin under the control of a β-estradiol-inducible promoter (LexOpr-*CLN1*). In the absence of β-estradiol, cells arrest in G1 due to the lack of expression of all G1 cyclins (*cln1*Δ *cln2*Δ *cln3*Δ) ^28,65^. This strain exhibited increased cell size following G1 arrest induction (Fig. S5A and B), and particle mobility increased with cell size expansion as expected (Fig. S5C and D). Deletion of *MPK1* further enhanced the cell volume expansion (Fig. S5A and B) and accelerated particle mobility during G1 arrest (Fig. S5C and D). These findings are fully consistent with our results using the *cdc28-as1* strain, suggesting that Mpk1-mediated control of cell size and cytoplasmic crowding is a general feature rather than a mutant-specific effect.

We next investigated which CWI signaling molecules are involved in maintaining cell size and cytoplasmic crowding during cell enlargement, besides Mpk1. We constructed *cdc28-as1* strains lacking each of the mechanosensory genes (*WSC1*, *WSC2*, *WSC3*, *MID2*, *MTL1*), MAPKKK gene (*BCK1*), and a transcriptional factor gene (*RLM1*) in the CWI pathway (Fig. 2D). In the absence of 1-NM-PP1 treatment, cell volumes and growth of these mutants were comparable to the parental strain *cdc28-as1* (Fig. S6A and B). After 1-NM-PP1 treatment, the cell volumes of *cdc28-as1* cells carrying *bck1*Δ or *mid2*Δ significantly increased to levels similar to those of *cdc28-as1 mpk1*Δ cells (Fig. 4E and F). Additionally, *cdc28-as1 wsc1*Δ and *cdc28-as1 rlm1*Δ cells exhibited a modest increase in cell volume relative to *cdc28-as1* cells, whereas *cdc28-as1* cells carrying *wsc2*Δ, *wsc3*Δ, or *mtl1*Δ did not exhibit significant changes (Fig. 4F). These results are consistent with the finding that the upstream components are required for Mpk1 phosphorylation, a well-established marker of Mpk1 activation (Fig. 2C). Next, we evaluated cytoplasmic crowding using 40nm-GEMs in these mutants. The *cdc28-as1* cells carrying *bck1*Δ, *mid2*Δ, or *wsc1*Δ exhibited higher median effective diffusion coefficients at 8 h after 1-NM-PP1 treatment than *cdc28-as1* cells (Fig. 4G). These results indicate that the core CWI components contribute to the regulation of cell size and cytoplasmic crowding during excessive growth.

### Mpk1 promotes cell wall thickening in enlarged cells

The cell wall constrains cell volume expansion by providing mechanical resistance to turgor pressure ^42^. We therefore asked whether disruption of the cell wall integrity perturbs the balance between turgor pressure and wall resistance. To address this, we examined and compared cell wall thickness in *cdc28-as1* and *cdc28-as1 mpk1*Δ cells using transmission electron microscopy (TEM) (Fig. 5A). Consistent with cell volume measurements (Fig. 4B), *cdc28-as1 mpk1*Δ cells were larger than *cdc28-as1* cells when treated with 1-NM-PP1 for 8 h (Fig. 5A). Interestingly, we found that *cdc28-as1* cells exhibited thicker cell walls than wild-type cells (Fig. 5B and C, S7A and B), suggesting the existence of mechanisms that promote cell wall thickening during excessive growth. Conversely, this cell wall thickening was not observed in *cdc28-as1 mpk1*Δ cells (Fig. 5B and C, S7B and C). The apparent electron density of the cell wall was also reduced in *cdc28-as1 mpk1*Δ cells compared with wild-type and *cdc28-as1* cells (Fig. 5B, S7A-C). Together, these results demonstrate that cells reinforce the cell wall during excessive growth, a process largely dependent on Mpk1.

**Fig. 5.**
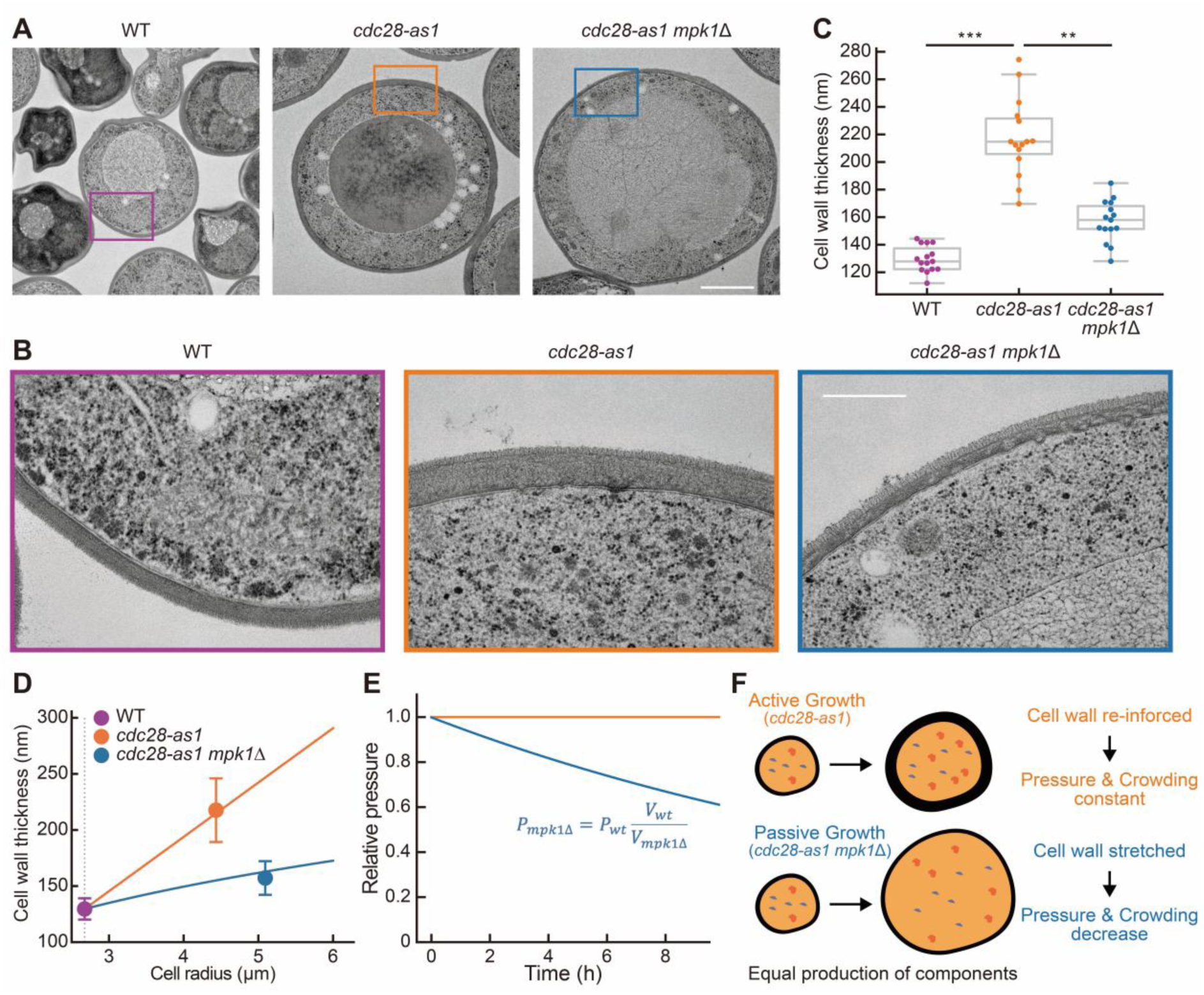
Measurement of cell wall thickness in enlarged cells. (A and B) Representative TEM images of wild-type (WT), *cdc28-as1*, and *cdc28-as1 mpk1*Δ strains. Cells were treated with 1-NM-PP1 for 8 h. (A) Scale bar, 2 μm. (B) Magnified images of the regions outlined by the square in panel A. Scale bar, 500 nm. (C) Quantification of cell wall thickness based on TEM images (n = 15 cells for each condition). Statistical significance was assessed using Kruskal-Wallis test followed by a *post-hoc* analysis by Dunn’s test with the Holm-Bonferroni correction. **, p < 0.01; ***, p < 0.001. In the boxplots, the center lines indicate the median, boxes represent the interquartile range, and whiskers extend to the minimum and maximum values, excluding outliers defined as 1.5 times the interquartile range. (D) Measured cell wall thickness from (C) and predicted cell wall thickness under two growth regimes based on Laplace’s law. Orange line: predicted values assuming constant turgor pressure and cell wall tension. Blue line: predicted values assuming passive stretching of the cell wall in *cdc28-as1 mpk1*Δ cells, leading to an excessive cell volume increase accompanied by a proportional decrease in turgor pressure. (E) Relative intracellular pressure derived from the measured volume expansions of *cdc28-as1* and *cdc28-as1 mpk1*Δcells. (F) Schematic models of active and passive growth illustrating the differences in macromolecular crowding and cell wall thickness between *cdc28-as1* and *cdc28-as1 mpk1*Δ cells.

### Cell wall thickening follows Laplace’s law for constant turgor and wall tension

Cell wall thickness is linked to cell volume, turgor pressure, and wall tension through Laplace’s law (see Materials and methods). If turgor pressure remains constant as cells grow, an increase in cell radius results in a proportional increase in tangential stress. Cell wall thickening can prevent excessive tensile stress in the cell wall, reducing the risk of mechanical failure. Strikingly, the measured cell wall thickness in enlarged *cdc28-as1* cells closely matched the theoretically predicted value that is required to maintain turgor pressure and the cell wall tension (Fig. 5D). In contrast, the thinner cell wall observed in the *cdc28-as1 mpk1*Δ enlarged cells does not follow this trajectory. Mpk1-dependent cell wall thickening therefore acts as a critical compensatory mechanism, reinforcing wall strength to maintain constant tensile stress and turgor pressure during cell enlargement.

Despite the differences in cell wall thickness between *cdc28-as1* and *cdc28-as1 mpk1*Δ cells, the total cell wall volume was nearly identical (Fig. S8A), suggesting that the CWI pathway promotes wall thickening not by increasing the synthesis of cell wall components but by preventing cell wall stretching. While the ratio of cell wall volume to total cell volume remains constant as *cdc28-as1* cells grow large, this ratio is reduced in *cdc28-as1 mpk1*Δ cells (Fig. S8B), indicating that cell volume increases disproportionately relative to cell wall synthesis in the absence of Mpk1. If the overall production rates of osmolytes and macromolecules remain unchanged in *cdc28-as1 mpk1*Δ cells, the accelerated volume expansion in these cells would reduce the internal pressure and dilute concentration of macromolecules. The predicted pressure decline resulting from this excess volume expansion would scale proportionally with the ratio of the cell volume between *cdc28-as1 mpk1*Δ and *cdc28-as1* cells (Fig. S8C). In that scenario, the pressure in *cdc28-as1 mpk1*Δ cells after 8 h of 1-NM-PP1 treatment would be 37% lower than that in *cdc28-as1* cells (Fig. 5E). Using this pressure, we also predicted the cell wall thickness required to maintain constant cell wall tension in *cdc28-as1 mpk1*Δ cells (Fig. 5D), which closely matched the measured cell wall thickness in these cells. Together, our data indicate that the CWI pathway regulates cell wall thickness in *cdc28-as1* cells to maintain cell wall tension and internal pressure constant during active growth (Fig. 5F). In the absence of this pathway, the cell wall fails to withstand the increased tangential stress in enlarged cells, leading to accelerated expansion and reduced cytoplasmic crowding in *cdc28-as1 mpk1*Δ cells, characteristic of passive growth (Fig. 5F).

### Loss of Mpk1 increases the sensitivity of enlarged cells to cell wall-related stresses

Deletion of Mpk1 leads to a thinner cell wall in *cdc28-as1* cells (Fig. 5C), which may reduce the mechanical resistance of the cell wall to external stresses. We therefore challenged cells with cell wall stresses using two approaches: (1) treatment with the cell wall lytic enzyme Zymolyase and (2) application of physical cell wall stress by sonication. Upon treatment with Zymolyase, we classified the responses of enlarged *cdc28-as1 mpk1*Δ cells into four categories: (1) burst, (2) blebbing, (3) spheroplast, and (4) normal (Fig. 6A and Movie S3). In this study, “burst” was defined as rupture of both the plasma membrane and the cell wall, resulting in leakage of intracellular contents into the extracellular space (Fig. 6A and B). “Blebbing” was defined as a transient protrusion of the plasma membrane caused by a local break in the cell wall, without ultimate disruption of the plasma membrane (Fig. 6A and B). “Spheroplast” was defined as the extrusion of the entire cell into the extracellular space after cell wall rupture, while the plasma membrane remains intact (Fig. 6A and B). All other cells were classified as “normal”. We prepared four samples by incubating *cdc28-as1* or *cdc28-as1 mpk1*Δ cells either with or without 1-NM-PP1 for 8 h. To evaluate the cell sensitivity to Zymolyase (1 U/mL), we first quantified the proportion of normal cells following the treatment (Fig. 6C). Quantification analysis revealed that enlarged *cdc28-as1 mpk1*Δ cells exhibited a sharp decline over time (Fig. 6C, blue). Enlarged *cdc28-as1* cells also showed a gradual decrease (Fig. 6C, orange), whereas no obvious decrease was detected in normally proliferating cells (Fig. 6C, gray and purple). After the Zymolyase treatment, the proportion of normal cells decreased to ∼20% in enlarged *cdc28-as1 mpk1*Δ cells and ∼80% in enlarged *cdc28-as1* cells, whereas the proportion remained close to 100% in normally proliferating cells without Zymolyase treatment (Fig. 6D). A similar overall trend was observed at a higher concentration of Zymolyase (10 U/mL), with the stronger effect characterized by a more rapid decrease in the number of normal cells (Fig. S9A and B). These results support the idea that Mpk1 is required for maintaining cell wall mechanical resistance during cell enlargement. We further classified cellular responses into four categories (Fig. 6A and B). Interestingly, approximately 20% of enlarged *cdc28-as1 mpk1*Δ cells exhibited blebbing at a lower concentration of Zymolyase (1 U/mL) (Fig. 6E). Similarly, at a higher concentration (10 U/mL), approximately 20% of enlarged *cdc28-as1 mpk1*Δ cells exhibited blebbing or spheroplast, whereas almost all cells underwent bursting in other samples (Fig. S9C). These blebbing and spheroplast cells were likely to lack sufficient internal pressure to rupture the plasma membrane, consistent with the Laplace’s law prediction of reduced turgor pressure in *cdc28-as1 mpk1*Δ cells (Fig. 5E).

**Fig. 6.**
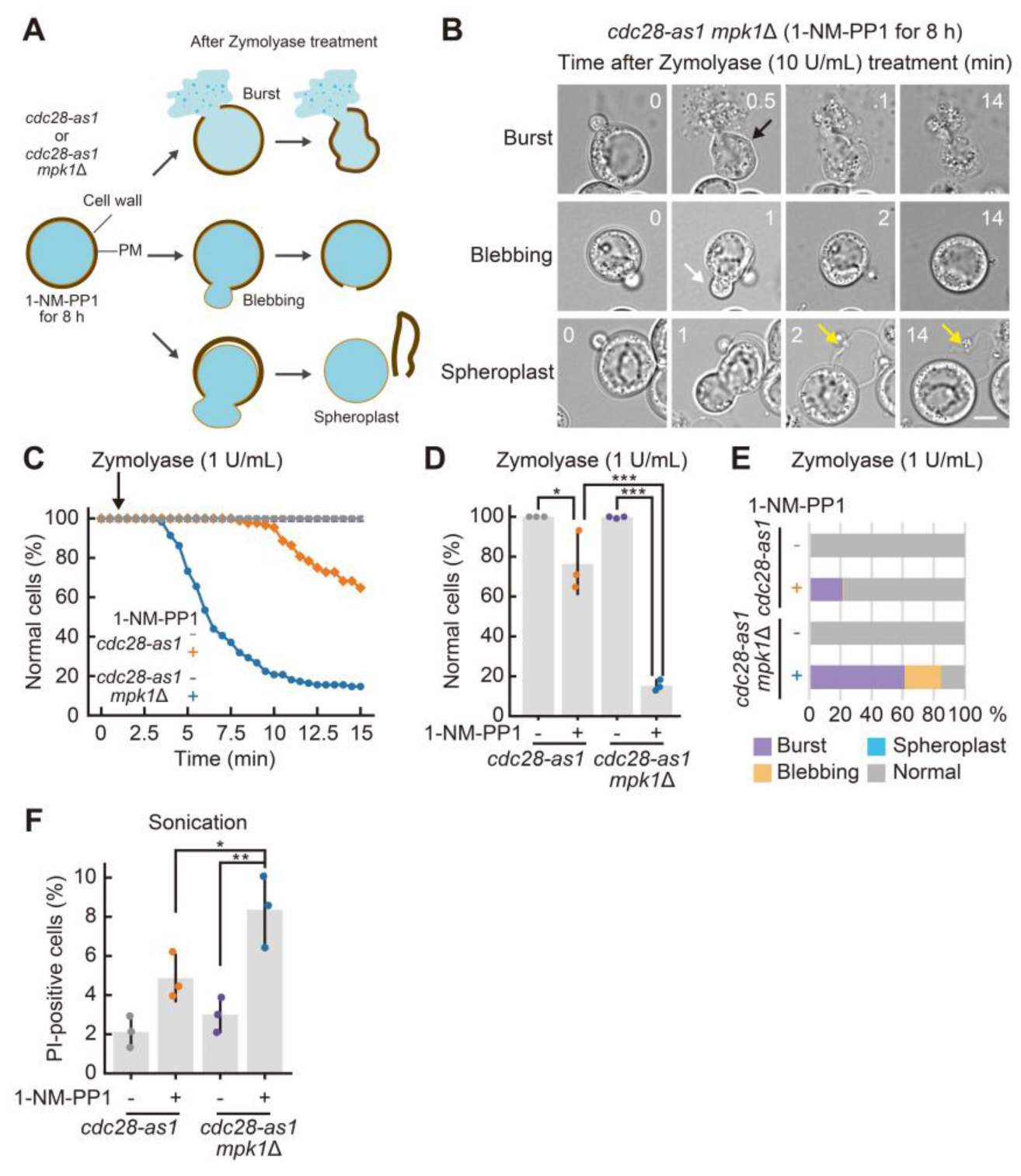
Loss of Mpk1 renders enlarged cells sensitive to cell wall stress. (A) A schematic illustration of the Zymolyase treatment experiments. PM; plasma membrane. (B) Representative time-lapse images of *cdc28-as1 mpk1*Δ cells treated with 1-NM-PP1 for 8 h followed by Zymolyase treatment (10 U/mL). The black arrow indicates the timing of bursting, the white arrow indicates blebbing, and yellow arrows indicate the cell wall. Scale bar, 5 μm. (C) Proportion of normal cells after Zymolyase treatment (1 U/mL) during 15 min time course (n > 86 cells). *cdc28-as1* without 1-NM-PP1: gray, *cdc28-as1* with 1-NM-PP1: orange, *cdc28-as1 mpk1*Δ without 1-NM-PP1: purple, and *cdc28-as1 mpk1*Δ with 1-NM-PP1: blue. (D) Proportion of normal cells at 14 min after Zymolyase treatment (1 U/mL). The mean values from the three independent experiments were represented with SD, and each dot represents the value from an individual experiment. (E) Classification of cells after 14 min of Zymolyase treatment (1 U/mL). Cells were categorized into four types: burst, blebbing, spheroplast, and normal (n > 250 cells from three independent measurements). (F) Difference in the proportion of PI-positive cells before and after sonication. The mean values from the three independent experiments were represented with SD, and each dot represents the value from an individual experiment (n > 130 cells for each experiment). (D and F) Statistical significance was assessed using one-way ANOVA followed by a *post hoc* Tukey-Kramer test. *, p < 0.05; **, p < 0.01; ***, p < 0.001.

Finally, we sonicated cells to impose physical cell wall stress. Cells were stained with PI before and after sonication to measure the fraction of dead cells, as PI-stained cells are considered as dead. Consistent with the results obtained with Zymolyase treatment (Fig. 6D and S9B), enlarged *cdc28-as1 mpk1*Δ cells exhibited the highest proportion of PI-positive cells amongst all samples (Fig. 6F). Taken together, these results indicate that Mpk1 is required to maintain cell wall homeostasis in response to cell wall stresses during excessive growth.

### Enlarged cells require Mpk1 for their survival after the cell cycle restart

A previous study reported that most yeast cells retain proliferative potential upon cell cycle restart following cell enlargement ^18^. However, accelerated cell enlargement and reduced cytoplasmic crowding caused by the loss of Mpk1 may lead to a decline in this potential. To test this possibility, cells were treated with 1-NM-PP1 for 8 h to generate enlarged cells and then released from the cell cycle arrest (hereafter referred to as cell cycle restart) (Fig. 7A). Enlarged *cdc28-as1* cells formed new buds after cell cycle restart, which differed from normal budding, as multiple buds were formed without cell separation (Fig. 7B and Movie S4). In contrast, enlarged *cdc28-as1 mpk1*Δ cells initially grew and formed a new bud; however, they subsequently underwent mother cell bursting within a few hours, likely due to rupture of the cell wall and plasma membrane (Fig. 7B and Movie S4). The proportion of non-burst cells gradually decreased over time in enlarged *cdc28-as1 mpk1*Δ cells, whereas no obvious decline was observed in enlarged *cdc28-as1* cells (Fig. 7C). At 12 h after cell cycle restart, only ∼15% of enlarged *cdc28-as1 mpk1*Δ cells remained non-burst, which was significantly lower than the proportion observed in *cdc28-as1* cells (< 90%) (Fig. 7D). Interestingly, CHX treatment partially rescued the decline in the proportion of non-burst cells in *cdc28-as1 mpk1*Δ cells (Fig. 7C), with ∼35% of cells remaining non-burst at 12 h (Fig. 7D). These results indicate that CWI pathway is required to prevent cell bursting upon cell cycle restart in excessively large cells.

**Fig. 7.**
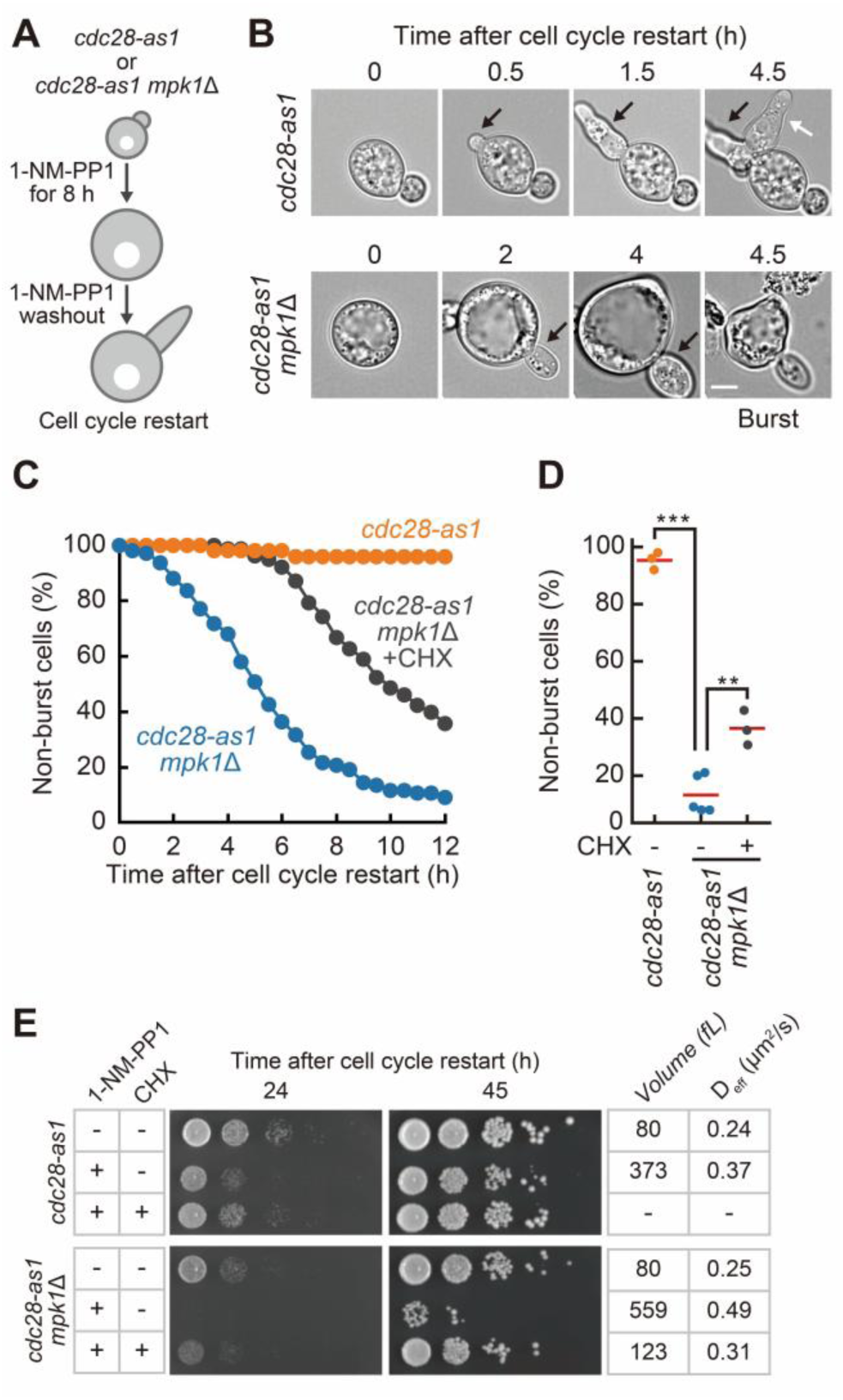
Enlarged cells require Mpk1 to survive upon the cell cycle restart. (A) A schematic illustration of the induction of cell cycle arrest and restart. *cdc28-as1* or *cdc28-as1 mpk1*Δ cells were incubated with 1-NM-PP1 for 8 h first to obtain enlarged cells. Cells were washed with media to release the cell cycle arrest, termed cell cycle restart. (B) Representative time-lapse images of *cdc28-as1* and *cdc28-as1 mpk1*Δ cells upon the cell cycle restart. Black and white arrows indicate the first and second budding events, respectively. Scale bar, 5 μm. (C) Proportion of non-burst cells after the cell cycle restart (*cdc28-as1*; n = 50, *cdc28-as1 mpk1*Δ; n = 110, *cdc28-as1 mpk1*Δ + CHX; n = 78 cells). (D) Proportion of non-burst cells at 12 h after the cell cycle restart. Red lines indicate mean values calculated from at least three independent measurements, and each dot represents the value from an individual experiment. Statistical significance was assessed using one-way ANOVA followed by a *post hoc* Tukey-Kramer test. **, p < 0.01; ***, p < 0.001. (E) *cdc28-as1* and *cdc28-as1 mpk1*Δ cells were cultured under three different cell culture conditions; no treatment, 1-NM-PP1 alone, or 1-NM-PP1+cycloheximide (CHX). After 8 h of incubation under each condition, cells were serially spotted onto a YPD plate at 10-fold dilution. Mean cell volume and median D_eff_ of 40nm-GEMs are indicated in the right table under each condition. The images were taken at 24 and 45 h after cell cycle restart.

Next, we examined the proliferative potential of enlarged cells using serial dilution assays. A delay in cell growth was detectable in enlarged *cdc28-as1* cells at 24 h following cell cycle restart, but a similar number of colonies were detectable at 45 h (Fig. 7E, lanes 1 and 2). By varying the duration of 1-NM-PP1 treatment to generate a range of cell sizes, we found that larger cells displayed more pronounced proliferation defects after 24 h recovery period, and cells eventually recovered proliferation after 45 h (Fig. S10A). These results indicate that cell enlargement slows down proliferation but most enlarged cells are able to form at least one viable daughter cell. In contrast, enlarged *cdc28-as1 mpk1*Δ cells formed fewer colonies compared with normally proliferating cells (Fig. 7E, lanes 4 and 5). The defect was partially rescued by CHX treatment (Fig. 7E, lanes 5 and 6), showing that loss of viability is a consequence of the increased size of these cells. Of note, we assessed cell viability by PI staining and found no detectable PI-positive cells in enlarged *cdc28-as1* and *cdc28-as1 mpk1*Δ strains before cell cycle restart, similar to normally proliferating cells (Fig. S10B), suggesting that nearly all enlarged cells were viable prior to restart. Furthermore, enlarged *cdc28-as1* cells carrying *bck1*Δ or *wsc1*Δ also failed to proliferate after cell cycle restart, similar to *cdc28-as1 mpk1*Δ cells (Fig. S10C). Together, these findings show that the CWI signaling pathway is required for efficient resumption of cell proliferation of enlarged cells following cell cycle restart.

## Discussion

Recent studies have shown that cytoplasmic biophysical properties change under diverse conditions and elucidated the mechanisms underlying their regulation in yeast cells ^9,13,15,16,66,67^. In this study, we demonstrate that cell volume expansion and cell wall remodeling need to be actively coordinated to maintain crowding homeostasis as cells grow larger. This regulation requires Mpk1 function, which becomes increasingly active as cell size increases. Mechanistically, Mpk1 ensures that the cell wall becomes thicker as cell volume expands, thereby providing the mechanical resistance required to counteract the increasing tangential forces generated by turgor pressure during cell enlargement. Our measurements of wall thickness match the values predicted by Laplace’s law. In the absence of Mpk1, cell walls are not reinforced sufficiently, leading to wall stretching, reduced crowding, and increased cell fragility, which precludes enlarged cells from successfully dividing.

The concentration of macromolecules inside cells is tightly controlled, and deviations from normal density coincide with altered cell function when cells differentiate, during cell senescence, or when cells respond to stress ^8,14–16,18^. It is widely accepted that changes in crowding are driven by an uncoupling of macromolecule biosynthesis and cell volume increase. Previous studies have shown that cell volume continues to expand even if macromolecule biosynthesis is attenuated, leading to cytoplasmic dilution in enlarged budding and fission yeast cells ^18,19,21,27^. Similarly, intracellular density in *S. pombe* and mammalian cells fluctuates during the cell cycle because volume expansion and biosynthesis are not tightly coupled throughout the cell cycle progression ^20,68–70^. Moreover, in *S. pombe*, transient inhibition of volume growth by osmotic oscillations while biosynthesis continues leads to an increase in intracellular density ^71^.

As water makes up more than 70% of the cell mass ^72^, cell volume is dictated by the osmotic pressure exerted by the cellular components. Macromolecule synthesis and volume increase can become uncoupled because osmotic pressure is dominated by small molecules, such as ions and metabolites, which are approximately 100-folds more concentrated than macromolecules in cells ^73,74^. In response to osmotic perturbations, cells restore cell volume and crowding homeostasis by modulating the concentration of small osmolytes ^75^. Our work uncovers a previously unrecognized mechanism of crowding control by altering the elastic properties of the cell wall. When cells grow larger, cell wall reinforcement is required to maintain crowding homeostasis (Fig. 5F). To support this, cell wall remodeling likely also contributes to density regulation in other conditions. In yeast spores, for example, the characteristic thick cell walls might help generate the densely crowded intracellular environment thought to protect spores against a broad range of external stresses ^13,66,76,77^.

We show that the CWI pathway is necessary to adjust cell wall thickness as cells grow larger (Fig. 5C). The CWI pathway controls the expression of the enzymes needed for the synthesis of the main cell wall components, glucan and chitin ^52,53^. However, our measurements show that the total volume of the cell wall remains unchanged regardless of CWI pathway activation (Fig. S8A), suggesting that the critical role of the CWI pathway is not to control the total amount of cell wall material, but rather to regulate its elastic properties. A critical determinant of cell wall stiffness is chitin, which makes up only a small fraction of the total cell wall mass yet strongly impacts its elastic properties ^78,79^. Whether increased chitin production is necessary to prevent cell wall stretching as cells grow larger remains to be determined.

Cells activate stress-response signaling pathways to halt cell cycle progression until underlying issue is resolved ^80,81^. These signaling pathways are often necessary for cell survival under the stress conditions. Our results show that the Mpk1 is necessary for cell cycle restart in enlarged cells, otherwise the enlarged cells burst (Fig. 7C and D). We therefore conclude that a key role of cell wall maintenance during cell enlargement is to preserve the proliferative potential in yeast. Our work highlights how signaling pathways evolved to ensure force balance as cell geometry changes. Similar mechanisms might exist in other species such that mammalian cells may employ cell wall independent mechanisms through cortical stiffening and/or cytoskeleton remodeling, that modulate plasma membrane tension ^82,83^. In plants, turgor pressure enforces mechanical demands on the cell wall, and dedicated signaling pathway monitor and adjust the cell wall properties during cell growth ^84,85^. More broadly, coupling of cell size sensing to cell cycle progression may exist as a conserved strategy to maintain cell size homeostasis in eukaryotes.

## Materials and methods

### Plasmids

All plasmids used in this study are listed in supplementary Table 1, along with Benchling links including the plasmid information, such as their sequences and maps.

### Budding yeast strains and culture conditions

All *S. cerevisiae* strains used in this study were haploid, derived from the W303 background, and are listed in supplementary Table 2 together with their origins. Yeast cells were basically grown in synthetic complete media (SC), composed of 2% glucose, 1% succinic acid, 0.6% sodium hydroxide, 0.5% ammonium sulfate, 1.7% yeast nitrogen base without amino acids and ammonium sulfate, with standard amino acid supplements. For liquid cultures, yeast cells were incubated in SC with gentle rotation. For serial dilution assays, yeast cells were grown on yeast extract peptone dextrose (YPD) agar plates, composed of 1% yeast extract, 2% peptone, 2% glucose, 60 μg/mL adenine, 20 μg/mL uracil, and 80 μg/mL tryptophan. Standard procedures were used for mating, tetrad analysis, and transformation.

To induce cell cycle arrest in *cdc28-as1* strains, cells were cultured in SC medium for 5–7 h at room temperature (25–27℃), diluted into fresh SC medium, and incubated overnight, ensuring that the optical density (OD_595_) did not exceed 0.5. Cells were then treated with 1-NM-PP1 (#TRC-A603003; LGC) at the final concentration of 10 μM and incubated at room temperature for the indicated duration. For release from cell cycle arrest, cells were collected by centrifugation, washed once with SC medium, and then resuspended in the fresh medium. To arrest the cell cycle in *cdc28-13* strains, cells were first grown in SC medium for 5–7 h at room temperature, diluted into fresh SC medium, and then incubated overnight to an OD_595_ below 0.5. Cells were shifted to the restrictive temperature 37℃ and incubated for the indicated duration. For LexOpr-*CLN1 cln1*Δ *cln2*Δ *cln3*Δ strains, cells were grown in SC medium supplemented with 10 nM β-estradiol (#E8875; Sigma-Aldrich) for 5–7 h at room temperature, diluted in the same medium, and incubated overnight. Cells were washed twice with SC medium to washout β-estradiol and incubated for the indicated duration to arrest cell cycle in G1.

### Serial dilution assays

Yeast cells were grown overnight in SC medium at room temperature to an OD_595_ of less than 1, diluted into fresh medium, and treated with 1-NM-PP1 at a final concentration of 10 μM when required. Cycloheximide (CHX; #C7698, Sigma-Aldrich) was also added to the medium at a final concentration of 1 μg/mL as needed. After incubation with gentle rotation at room temperature for the indicated duration, cells were collected by centrifugation, washed with SC medium, and resuspended in the same medium. Cells were then serially spotted at 10-fold dilution for 5 times on YPD plates, and grown at 30℃ for the indicated duration. The ODs were adjusted between samples before plating in Fig. S6B. Because differences in cell volume may affect turbidity, cell numbers, rather than ODs, were adjusted across samples in Fig. 7E, S10A, and S10C.

### Cell volume quantification

Yeast cells were sonicated for 1 second using a Misonix Sonicator XL-2000 (Qsonica) with lowest setting and then diluted in Isoton Ⅱ (7.93 mg/mL NaCl, 0.38 mg/mL disodium EDTA, 0.4 mg/mL KCl, 0.22 mg/mL NaH_2_PO_4_, 2.45 mg/mL Na_2_HPO_4_, and 0.3 mg/mL NaF). Mean cell volumes were determined using a Multisizer 3 Coulter counter (Beckman Coulter). The total number of particles was set to 60,000.

### Flow cytometric analysis of DNA content

Cells were collected by centrifugation, resuspended in 70% ethanol for fixation, and stored at 4℃ until use. Cells were sonicated for 10 seconds using a Misonix Sonicator XL-2000 with lowest setting, pelleted, and resuspended in 50 mM Tris-HCl buffer (pH 7.8). Cells were washed once with 50 mM Tris-HCl buffer (pH 7.8) and incubated with 2 mg/mL RNase A in the same buffer overnight at 37℃. Cells were treated with 5 mg/mL pepsin (#P7000; Sigma-Aldrich) for 30 min at 37℃, washed with Tris-HCl/NaCl/MgCl_2_ buffer (200 mM Tris-HCl (pH 7.5), 211 mM NaCl, and 78 mM MgCl_2_), and resuspended in the same buffer. Cells were stained with Propidium iodide (PI; #P4170, Sigma-Aldrich) at a final concentration of 75 μg/mL. DNA content was analyzed using a BD Accuri C6 flow cytometer (BD Biosciences). A total of 20,000 cells were analyzed per sample, and fluorescence signals were detected using the FL2 channel. To analyze DNA content from flow cytometer data, histograms were generated after excluding values other than the 1C and 2C populations.

### Transmission electron microscopy (TEM) observation

Yeast cells were grown in SC medium at room temperature for 5 h to an OD_595_ of 0.3–0.7. Cultures were diluted into fresh medium and grown overnight to an OD_595_ below 0.7. *cdc28-as1*, *cdc28-as1 mpk1*Δ, and wild-type strains were treated with 1-NM-PP1 at a final concentration of 10 μM, cultured for 8 h, and then harvested. Subsequent procedures were performed by Tokai Electron Microscopy, Inc. Yeast cells were sandwiched between copper plates and rapidly frozen in liquefied propane. Samples were subjected to freeze substitution in ethanol containing 2% glutaraldehyde and 1% tannic acid at −80 °C for 48 h, −20 °C for 3 h, and 4 °C for 3 h, followed by warming to room temperature. Cells were dehydrated for 30 min three times and overnight in anhydrous ethanol. Cells were immersed twice in propylene oxide for 30 min, then in a 1:1 mixture of propylene oxide and resin (Quetol-651; Nisshin EM Co.) for 3 h, and finally in resin overnight. Samples were embedded in resin at 60°C for 48 h. Embedded cells were cut with an ultramicrotome (Ultracut UCT; Leica) to prepare 70-nm-thick sections, stained with 2% uranyl acetate for 15 min and lead stain solution (Sigma-Aldrich) for 3 min, and observed with a transmission electron microscopy (JEM-1400 Plus; JEOL Ltd.). To measure cell wall thickness from TEM images, three magnified images were obtained per cell, and the mean thickness was calculated for each cell.

### Live-cell imaging

Yeast cells were imaged with an Ti2-E inverted microscope (Nikon) equipped with an sCMOS camera (ORCA-Fusion BT; Hamamatsu Photonics), an oil-immersion objective lens (CFI Plan Apochromat Lambda 60XC, NA = 0.95, WD = 0.21-0.11 mm; Nikon). The illumination light source was a SOLA Light Engine (Lumencor). The fluorescence filters were as follows: high signal-to-noise BL series filter cubes GFP/FITC/Cy2 (Nikon) (excitation filter, 466/40; dichroic mirror, 495; emission filter, 525/50) for 40nm-GEMs and mNeonGreen and Texas Red/mCherry (Nikon) (excitation filter, 562/40; dichroic mirror, 593; emission filter, 641/75) for mScarlet-I. Images were acquired using NIS-Elements Advanced Research (Nikon).

To image cells using 96-well glass-bottom plates (P96-1.5H-N; Cellvis), cells were transferred to the wells precoated with 1 mg/mL concanavalin A (#L7647; Sigma-Aldrich) and allowed to settle onto the glass surface for 5 min. Cells were then washed once with fresh medium and resuspended in the same medium. All fluorescence and time-lapse images were taken using 96-well glass-bottom plates.

For 40nm-GEMs imaging, the fluorescence images were recorded every 20 milliseconds for a total of 4 seconds. For Mpk1- or Htb1-mNeonGreen together with Nup49-mScarlet-I expressing cells, z-stacks spanning 8–12 µm were taken at 0.5-µm intervals to cover the entire cell. The z-stacks were processed using maximal intensity projection. To measure the budding index, cells were sonicated for 1 second using a Misonix Sonicator XL-2000 with lowest setting, concentrated by centrifugation, mounted on glass slides (thickness, 1.0 mm; Thermo Fisher Scientific), and sealed with glass coverslips (thickness, 0.16–0.19 mm; Northwest Scientific). Cells were imaged and classified as budded or unbudded based on visual inspection.

### Cell wall stress induced by Zymolyase and sonication

For the Zymolyase (#E1005; Zymo Research) treatment experiments, *cdc28-as1* and *cdc28-as1 mpk1*Δ strains were cultured in SC medium supplemented with 10 μM 1-NM-PP1 at room temperature for 8 h. Control experiments were performed without 1-NM-PP1 treatment. Cells were transferred to 96-well glass-bottom plates precoated with concanavalin A and allowed to settle down to the bottom for 5 min. Cells were then washed once with SC medium containing 10 μM 1-NM-PP1 and resuspended in the same medium. Time-lapse imaging was performed at 30-sec intervals. Zymolyase was added during imaging to a final concentration of either 1 or 10 U/mL after the first minute, and imaging was continued for a total duration of 15 min. For the analysis, cells were categorized into (1) burst, (2) blebbing, (3) spheroplast, and (4) normal based on visual inspection. To calculate the half-time required for bursting to reach a plateau following 10 U/mL Zymolyase treatment, the data were fitted with a logistic regression model using Python 3 (https://www.python.org/) and Scipy 1.10.1 (https://scipy.org/).

For the sonication treatment experiments, cells were prepared in the same manner as for the Zymolyase treatment. Cells were sonicated for 3 seconds using a Misonix Sonicator XL-2000 with lowest setting. As a control, cells without the sonication were also prepared. To stain cells with PI, cultures were transferred to 96-well glass-bottom plates in the presence of PI at a final concentration of 37.5 μg/mL. After 5 min, cells were imaged by a microscope. For the analysis, PI-positive cells were counted, and the percentage of PI-positive cells was calculated as the ratio of PI-positive cells to the total number of cells. Figure 6F showed values obtained by subtracting the measurements of untreated cells from those of sonicated cells.

### Particle trajectory analysis

The GEM particles were tracked using the Mosaic suite, a FIJI plugin ^86^. In the analysis using Mosaic suite, the following parameters were used: radius = 3; cutoff = 0; per/abs, variable; link range = 1; and maximum displacement = 6 pixels, assuming Brownian dynamics. Following the particle tracking using Mosaic suite, individual particle trajectories were analyzed with the GEMspa software package ^87^. To calculate the effective diffusion coefficient (D_eff_) of individual particles, mean-square displacement (MSD) was computed using trajectories containing more than 10 time points. MSD curves were truncated to the first 10 points (200 msec) and fitted with a linear relationship to obtain the time-averaged MSD according to the equation: MSD(*T*) = 4D_eff_*T*, where *T* denotes the imaging time interval and D_eff_ is expressed in units of μm^2^/s. Regions of interest (ROIs) corresponding to individual cells were generated from bright-field images using Cellpose3^88^. These ROIs were then imported into GEMspa to quantify effective diffusion at single-cell level. For single-cell analyses, the median D_eff_ value from all trajectories within each cell was used as a representative value. For time-course analyses, the median D_eff_ value from all trajectories at each time point was used. For particle trajectory analysis, data from cells containing aggregated particles or small buds were excluded.

### Sample preparation for phosphoproteomics analysis

*cdc28-13* yeast cells were streaked onto a YPD plate and incubated overnight at room temperature. Cells were then cultured at 25℃ in YPD and diluted to an OD_600_ of 0.002 before being incubated overnight at 25℃. To ensure entry into the exponential growth phase, the overnight cultures were diluted the next morning and grown at 25℃ until reaching an OD_600_ of 0.30. A 500 mL culture of *cdc28-13* cells was then incubated at the restrictive temperature of 37℃, with samples collected at 2, 4, and 6 h. TCA (final concentration 6.25%) was added to the samples followed by shaking and 10 min incubation. Pellets were collected by centrifugation (1,500 g, 5 min, 4℃), followed by washing with ice-cold acetone twice. Pelleted yeast was resuspended in an equal volume of lysis buffer (8 M urea, 100 mM ammonium bicarbonate, 5 mM EDTA, pH 8) and ice-cooled glass beads. Cells were lysed using a bead beater (6 m/s, 30 s) for 5 min, followed by centrifugation at maximum speed for 5 min at 4℃. The supernatant was collected, and the lysis process was repeated five times with fresh buffer added to the pellet each time. Supernatants were stored at - 20℃.

A total of 5 mg of protein (quantified via BCA analysis) was reduced with 5 mM tris (2-carboxyethyl) phosphine (TCEP) for 30 min, alkylated with 12 mM iodoacetamide for 30 min in the dark, and diluted to 1 M urea before digestion. Proteins were digested overnight at 37℃ using sequencing-grade trypsin (Promega) at a 1:100 (w/w) enzyme-to-protein ratio. The following day the proteolysis was quenched by adding 100% formic acid until pH 3 and peptides were subjected to C18 cleanup (C18 3cc Water SepPak columns).

Peptides were first loaded onto a column and subjected to sequential washes with 1 column volume (CV) of methanol followed by 1 CV of 50% acetonitrile. The column was then equilibrated with 0.1% formic acid. After equilibration, the column was washed with 3 CV of 0.1% formic acid, and peptides were eluted using 2 CV of 50% acetonitrile in 0.1% formic acid. The eluted peptides were dried in a vacuum centrifuge at 30℃ and resuspended in 20 µL of 50% acetonitrile containing 0.1% formic acid. For LC-MS analysis to normalize phosphorylation levels in non-enriched samples, 2 µL of the peptide solution was diluted to 5% acetonitrile and analyzed.

For phosphopeptide enrichment, the remaining peptides (18 µL) were diluted 1:10 in phospho-enrichment loading buffer, which consisted of 50% acetonitrile, 29.2% lactic acid, and 0.1% trifluoroacetic acid (TFA). Low-binding stage tips were prepared by inserting two C18 disks, which were pre-washed with methanol. TiO2 resin (GL Science) was loaded into the stage tips and equilibrated twice with 100 µL of phospho-enrichment loading buffer. The peptide mixture was then incubated with the TiO2 resin by passing it through the stage tips via centrifugation at 100 g.

The resin was washed sequentially with 100 µL of (1) phospho-enrichment loading buffer, (2) 80% acetonitrile containing 0.1% TFA, and (3) 50% acetonitrile containing 0.1% TFA. Phosphopeptides were eluted in 250 µL of 50 mM ammonium hydroxide (pH ∼10.5) followed by 30 µL of 50% acetonitrile containing 0.1% formic acid. The eluted peptides were neutralized by dilution in 100 µL of 3% formic acid.

To desalt the phosphopeptides, C18 stage tips were prepared following the protocol described by Rappsilber et al. ^89^. Briefly, C18 disks were washed with methanol and 50% acetonitrile, then equilibrated with 0.1% formic acid before loading the phosphopeptides. After washing with 0.1% formic acid, the phosphopeptides were eluted in 100 µL of 50% acetonitrile containing 0.1% formic acid.

### Phosphoproteomics data acquisition

LC-MS/MS analysis was performed on an Orbitrap Q Exactive Plus mass spectrometer (Thermo Scientific) coupled to an EASY-nLC-1000 system (Thermo Scientific). Peptides were separated on an Acclaim PepMap 100 C18 (25 cm length, 75 µm inner diameter) with a two-step linear gradient from 3% to 25% acetonitrile in 110 min and from 25% to 40% acetonitrile in 10 min at a flow rate of 300 nL/min.

The DDA acquisition mode for phospho dataset was set to perform one MS1 scan followed by 20 MS2 scans (TOP20). The MS1 scan was performed in the Orbitrap (R = 70,000, 3e6 AGC target, maximum injection time of 64 ms and scan range 350–1500 m/z). Peptides with charge state between 2–6 were selected for fragmentation (isolation window: 1.4 m/z and fragmentation with HCD, intensity threshold 1.8e4, NCE 25%) and MS2 scans were acquired in a Orbitrap (R = 35,000, 5e5 AGC target, maximum injection time of 110 ms and scan range 200 to 2,000). A dynamic exclusion of 30 s was applied.

The DDA acquisition mode for proteome profiling was set to perform one MS1 scan followed by 20 MS2 scans (TOP20). The MS1 scan was performed in the Orbitrap (R = 70,000, 3e6 AGC target, maximum injection time of 64 ms and scan range 350–1500 m/z). Peptides with charge state between 2–6 were selected for fragmentation (isolation window: 1.4 m/z and fragmentation with HCD, intensity threshold 3.6e4, NCE 25%) and MS2 scans were acquired in a Orbitrap (R = 17,500, 1e5 AGC target, maximum injection time of 55 ms and scan range 200 to 2,000). A dynamic exclusion of 30 s was applied.

### Phosphoproteomics data analysis

MS spectra were analyzed using the MaxQuant software package (version 1.5.2.8) ^90^. Spectra were searched against the *S. cerevisiae* protein database downloaded from UniProt (May 2021) containing 6,055 entries, supplemented with common contaminants. Enzyme specificity was set to Trypsin/P allowing up to two missed cleavages. Carbamidomethylation of cysteine residues was set as a fixed modification, oxidation of methionine and deamidation of asparagine and phosphorylation of serine, threonine and tyrosine (this last modification only for the phosphor dataset) were specified as variable modifications. The first search peptide mass tolerance was set to 10 ppm and the main search peptide mass tolerance to 4.5 ppm. MS/MS fragment ion tolerance was set to 20 ppm for Orbitrap measurements. The minimum peptide length was set to seven amino acids and the maximum number of variable modifications per peptide was limited to three. Label-free quantification (LFQ) was enabled with a minimum ratio count of two, “match between runs” feature was disabled. Peptide and protein identifications were filtered using a target–decoy approach to achieve a false discovery rate (FDR) of 1% at both the peptide-spectrum match and protein levels.

The “Phospho (STY) Sites.txt” output table by was imported and filtered to remove reverse database hits and common contaminants. Only phosphosites with a posterior error probability (PEP) < 0.01 and localization probability > 0.75 were retained. Sites detected in at least two replicates within at least one experimental condition were kept for downstream analysis, resulting in 7,743 quantified phosphosites. Data were log_2_-transformed and median normalized across samples. Missing values were imputed in two steps: first, when a phosphosite was missing in only one replicate within a condition, the value was replaced by the mean of the remaining replicates; second, remaining missing values (∼8% of the data matrix) were imputed by random sampling from a normal distribution derived from the lowest 1% intensity values in the dataset.

For the proteome abundance “proteinGroups.txt” output table was imported and filtered to remove reverse database hits and common contaminants. Only proteins identified with at maximum one missing value per condition were considered, resulting in the identification of 3,356 proteins. Imputation of missing values was performed with a two step as described for the phosphor dataset.

Phosphosite intensities were normalized to the corresponding protein abundance to account for protein-level changes (4,985 phosphosites). Differential phosphorylation levels were assessed using one-way analysis of variance (ANOVA) and Tukey’s post hoc test with FDR using Benjamini–Hochberg. Phosphosites showing significant differences between at least one pair of conditions (adjusted p-value adjusted < 0.05) were considered regulated and retained for cluster analyses (1,548 phosphosites). For clustering analysis, phosphosite intensities were converted to Z-scores across samples. Time-resolved phosphorylation patterns were identified using fuzzy c-means clustering implemented in the Mfuzz R package ^91^. Cluster stability analysis was performed to define the number of clusters (8) Phosphosites with a cluster membership value ≥ 0.6 were considered core members of each cluster. Predicted upstream kinases were identified using the NetworKIN algorithm ^49^. Only kinase predictions with a score > 2 were considered. Kinase enrichment within each phosphorylation cluster was assessed using a hypergeometric test, comparing the number of predicted kinase targets within each cluster to their frequency across the full phosphosite dataset ^48^.

For the proteome analysis, differential protein levels were assessed using one-way analysis of variance (ANOVA) and Tukey’s post hoc test with FDR using Benjamini–Hochberg.

### Derivation of the cell wall model based on Laplace’s law

We model a cell as a growing sphere. Let *V*(*t*) be the cell volume and *R*(*t*) be the cell radius at time *t*. Both *cdc28-as1* and *cdc28-as1 mpk1*Δ are observed to grow exponentially over time:

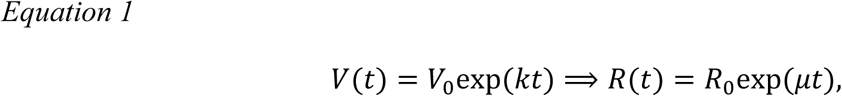

where *k* is the volumetric growth rate and *μ* = *k*/3 is the radial growth rate, which is different for each strain, and given by fitting the experimental data (Fig. S8C). At *t* = 0, both strains share the same initial radius *R*_0_ and initial wall thickness *w*_0_. The mechanical tension within the cell wall is given by Laplace’s law:

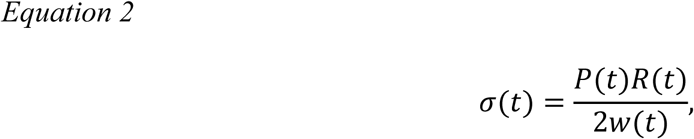

where *σ*(*t*) is the tensile stress, *P*(*t*) is the internal turgor pressure, and *w*(*t*) is the thickness of the cell wall.

#### Active growth

Cells need to prevent excessive stress in the cell wall and keep the concentration of small and large molecules constant. To achieve homeostasis during growth, cells therefore need to maintain the internal turgor pressure and the cell wall stress constant, meaning *P*(*t*) = *P*_0_ and *σ*(*t*) = *σ*_0_. In this case, Laplace’s law is written as:

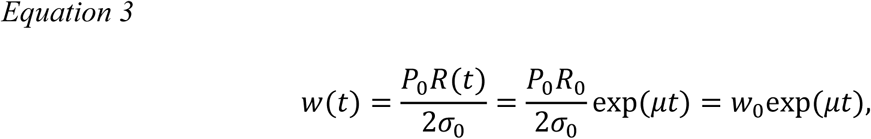

or

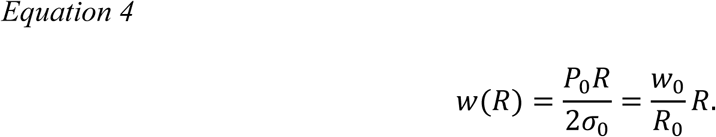

To achieve homeostasis during growth in this case, cells need to thicken their cell wall and Laplace’s law lets us predict the necessary cell wall thickness to achieve this. Since *w*_0_ and *R*_0_are given by the experimental data, we can predict *w*(*R*). Figure 4D shows that the cell wall thickness measured in arrested *cdc28-as1* cells aligns with this prediction. Cells are therefore able to adjust their cell wall thickness sufficiently to allow the cell wall tension and pressure to remain constant during growth. In contrast, the thinner cell wall observed in the larger arrested *cdc28-as1 mpk1*Δ indicates that cells are not able to achieve pressure and tension homeostasis. The observed thinning of the cell wall at larger size is only possible if either the pressure is decreased or the tension in the wall increased.

#### Passive growth

Experimental data in Figure 4D and Figure S8C shows that while the cell volume and cell wall thickness differ between *cdc28-as1* and *cdc28-as1 mpk1*Δ cells, the total cell wall volume is the same (Fig. S8A), suggesting that the production of biomass continues at the same rate in *cdc28-as1* and *cdc28-as1 mpk1*Δ cells. This indicates that the CWI pathway promotes thickening of the wall by preventing it from being stretched, rather than by promoting the synthesis of cell wall material. We therefore wondered how the turgor pressure would change if *cdc28-as1 mpk1*Δ cells continue to produce macromolecules and osmolytes at the same rate as wild type cells, but expand at the rate we observed in the data. In this situation, the total number of solute molecules *n*(*t*) is identical in both cell types: *n*_*mpk*1Δ_(*t*) = *n*_*wt*_(*t*). We will use the notation *mpk*1Δ to refer to the *cdc28-as1 mpk1*Δ cells, and *wt* to refer to the *cdc28-as1* cells. Because the *cdc28-as1 mpk1*Δ cells expand their volume at a faster rate (*k*_*mpk*1Δ_ > *k*_*wt*_), these solutes are being diluted when expanding. Using van ’t Hoff’s equation, *P* ∝ *n*/*V*, the internal pressure of the *cdc28-as1 mpk1*Δ cells decay relative to the *cdc28-as1* as:

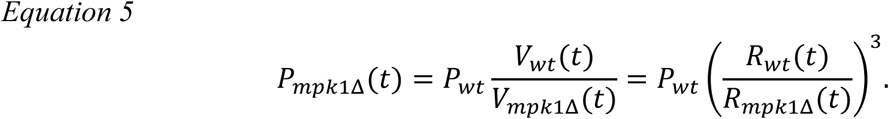

Substituting the pressure scaling *P*_*mpk*1Δ_(*t*) into Laplace’s law yields:

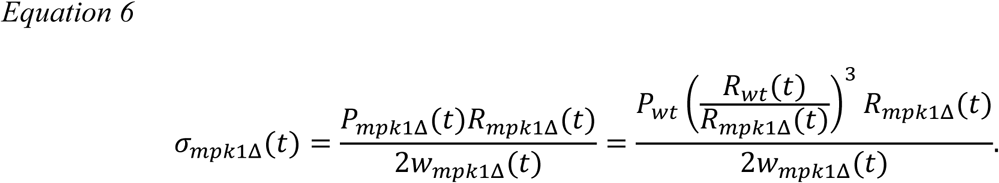

We assume that *cdc28-as1 mpk1*Δ cells still aim to maintain the same safe mechanical stress as the initial WT state 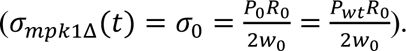 Therefore, canceling common terms (*P_wt_* and 2) leads to:

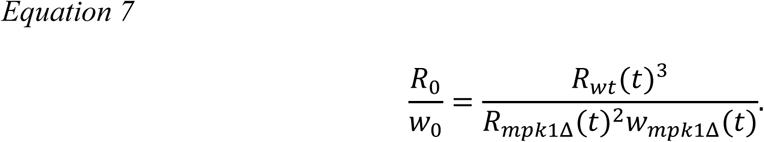

Solving for the wall thickness:

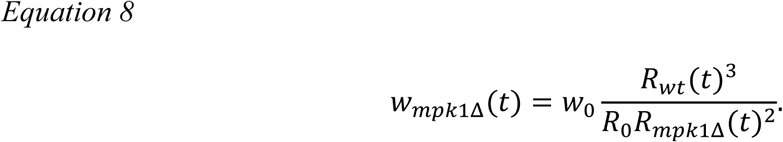

We then define the dimensionless ratio *α*, as

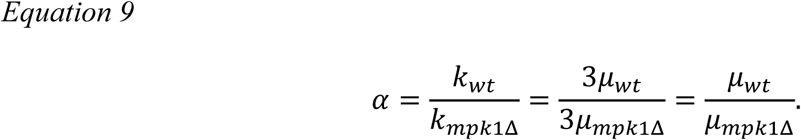

By isolating time *t* from the radial growth equation 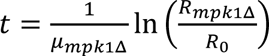 and substituting it into the growth equation, we can express *R*_*wt*_ strictly as a function of *R*_*mpk*1Δ_:

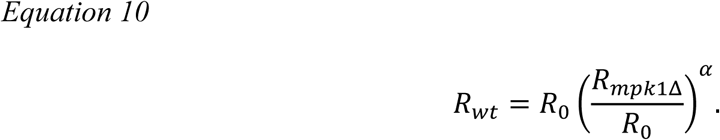

Substituting this relation leads to:

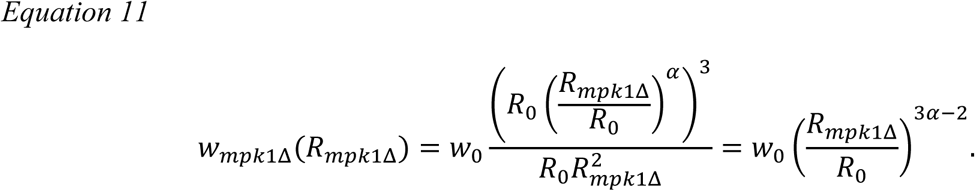

We can then predict the thickness of the wall for this passive growth regime using equation 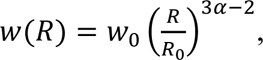 with *w*_0_, *R*_0_ and *α* determined from the experimental growth data. This allows us to calculate how the thickness should vary if the cell is growing passively at the measured rate. Figure 4D shows that the cell wall thickness measured in arrested *cdc28-as1 mpk1*Δ cells align with this prediction. This shows that the decrease of turgor pressure that results from the disproportionally fast volume expansion in *cdc28-as1 mpk1*Δ is sufficient to explain the altered cell wall thickness in these cells.

### Images and data analysis

All fluorescence imaging data were analyzed and quantified using Fiji/Image J (https://fiji.sc/). Background was subtracted on all fluorescence images using the rolling ball method (radius, 50.0 pixels). Data visualization and graph generation were performed using Python 3.11.4 with the Numpy 1.24.4, Pandas 1.5.3, Matplotlib 3.7.1, and Seaborn 0.13.2 modules.

For Figure 3 and S3, images of Mpk1- or Htb1-mNeonGreen and Nup49-mScarlet-I were obtained as 17–25 z-slices at 0.5 µm intervals and shown as two-dimensional images by the maximal intensity projection, as indicated in the figure legends. Whole cells and nuclei were segmented using Mpk1-mNeonGreen and Nup49-mScarlet-I images, respectively, with Cellpose3 ^88^, and fluorescence intensities of Mpk1- or Htb1-mNeonGreen were quantified. Cell area was measured from Mpk1-mNeonGreen images with Cellpose3 and used as a proxy for cell size.

Cells expressing 40nm-GEMs were segmented using bright-field images with Cellpose3, and cell area was measured as a proxy for cell size. From the image series (201 frames) of 40nm-GEMs, the first frame was shown as a representative image.

## Abbreviations

CWI: Cell wall integrity
MAPK: Mitogen-activated protein kinase
MAPKKK: MAP kinase kinase kinase
GEMs: Genetically encoded multimeric nanoparticles
D_eff_: Effective diffusion coefficient
MSD: Mean-square displacement
mNG: mNeonGreen
CHX: Cycloheximide
PI: Propidium iodide

## Acknowledgments

We would like to thank all members of the Ikui laboratory for their helpful discussions and technical assistance. We thank Lea Goddard for technical support. Some budding yeast strains were kind gifts from Dr. David O. Morgan, Dr. Frederick R. Cross, Dr. Jan M. Skotheim, and Dr. Matthew P. Swaffer. Some plasmids were kind gifts from Dr. Liam J. Holt and we would like to thank Dr. Ying Xie for kind support with handling these plasmids. We are grateful to Dr. Yohei Kondo for his valuable advice on data analysis. We are also grateful to the late Dr. Angelika Amon for bringing A.I. and G.E.N. together. This work was supported by National Institute of Health (SC1GM121242) and NSF (2508688) to A.I.. G.E.N. was supported by an SNSF Project Funding Grant (310030_212660). K.S. was supported by The Uehara Memorial Foundation.

## Competing Interests

No competing interests declared.

## Author contributions

Conceptualization: K.S., G.E.N., A.I.; Data curation: K.S., M.K., F.U., L.W., R.L., Y.K.; Formal analysis: K.S., M.K., F.U., L.W., R.L.; Funding acquisition: G.E.N., A.I.; Investigation: K.S., M.K., F.U., L.W., R.L., Y.K.; Methodology: K.S., M.K., F.U., L.W., R.L., Y.K.; Project administration: G.E.N., A.I.; Resources: K.S., M.K., F.U., L.W., R.L.; Software: F.U., L.W.; Supervision: G.E.N., A.I.; Validation: K.S., M.K., F.U., L.W., R.L.; Visualization: K.S., M.K., F.U., L.W., R.L.; Writing – original draft: K.S., M.K., F.U., L.W.; Writing - Review & Editing: K.S., M.K., F.U., L.W., Y.K., G.E.N., A.I.

## Notes

### Competing Interest Statement

The authors have declared no competing interest.

