## Supplemental File 1 for "Cell wall remodeling is required for crowding homeostasis during cell growth in *S. cerevisiae*"

excessive cell growth, cytoplasmic crowding, cell wall remodeling, CWI signaling pathway, MAPK, budding yeast

**Abbreviations:**

| | |
|---|---|
| CWI | Cell wall integrity |
| MAPK | Mitogen-activated protein kinase |
| MAPKKK | MAP kinase kinase kinase |
| GEMs | Genetically encoded multimeric nanoparticles |
| $D_{eff}$ | Effective diffusion coefficient |
| MSD | Mean-square displacement |
| mNG | mNeonGreen |
| CHX | Cycloheximide |
| PI | Propidium iodide |

**Supplementary information**

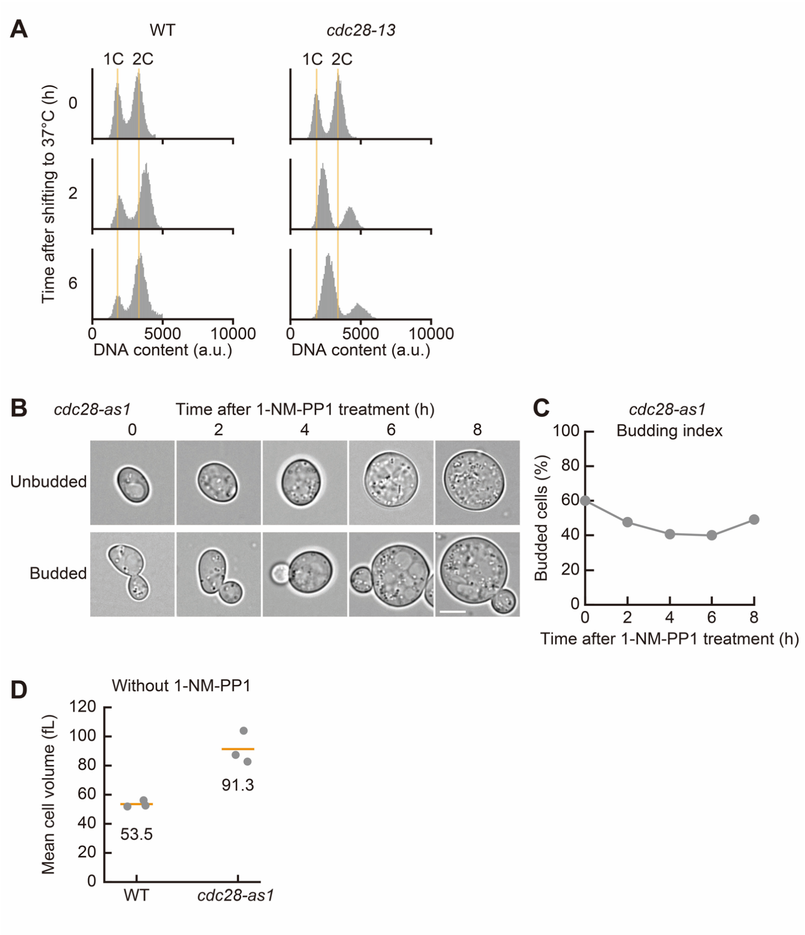


**Fig. S1. Characterization of the *cdc28-13* and *cdc28-as1* cells after cell cycle arrest induction.**

(A) DNA content determined by flow cytometry in wild-type (WT) and *cdc28-13* cells after shifting to the restrictive temperature 37°C.

(B) Representative bright field images of unbudded (upper) and budded (lower) *cdc28-as1* cells after 1-NM-PP1 treatment. Scale bar, 5 µm.

(C) Quantification of budding index in *cdc28-as1* cells after 1-NM-PP1 treatment (n > 100 cells).

(D) Cell volume determined using a Coulter counter in WT and *cdc28-as1* cells in the absence of 1-NM-PP1. Orange lines indicate the mean values calculated from three independent measurements, and each dot represents the value obtained from an individual experiment. The indicated values correspond to the mean cell volumes.

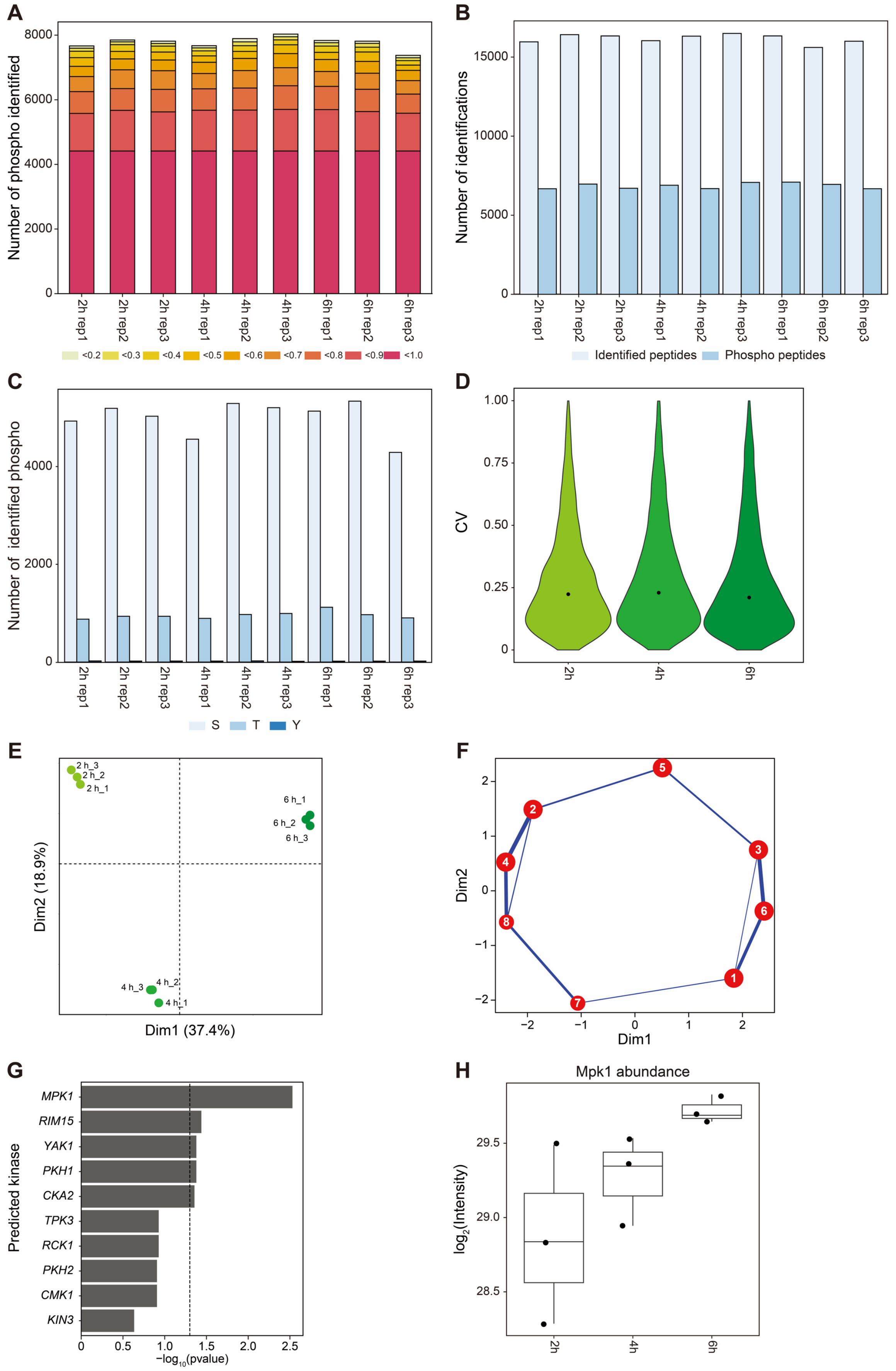

A
Number of phospho identified
8000
6000
4000
2000
0
2h rep1
2h rep2
2h rep3
4h rep1
4h rep2
4h rep3
6h rep1
6h rep2
6h rep3
<0.2 <0.3 <0.4 <0.5 <0.6 <0.7 <0.8 <0.9 <1.0
B
Number of identifications
15000
10000
5000
0
Identified peptides
Phospho peptides
C
Number of identified phospho
4000
2000
0
S
T
Y
D
CV
1.00
0.75
0.50
0.25
0
2h
4h
6h
E
2 h_3
2 h_2
2 h_1
6 h_1
6 h_2
6 h_3
4 h_3
4 h_2
4 h_1
Dim2 (18.9%)
Dim1 (37.4%)
F
Dim2
Dim1
G
Predicted kinase
MPK1
RIM15
YAK1
PKH1
CKA2
TPK3
RCK1
PKH2
CMK1
KIN3
0
0.5
1.0
1.5
2.0
2.5
−log10(pvalue)
H
Mpk1 abundance
log2(Intensity)
29.5
29.0
28.5
2h
4h
6h

**Fig. S2. Quality control of the phosphoproteomics dataset.**

(A) Number of identified phosphosites across all analyzed conditions. The completeness of phosphosite identification across samples is indicated by a color scale: dark red represents phosphosites identified in all conditions, whereas pale yellow indicates phosphosites detected only sparsely across the dataset.

(B) Phosphorylation enrichment across the analyzed conditions. The total number of identified peptides and phosphopeptides is shown for each sample in light blue and dark blue, respectively.

(C) Distribution of phosphorylated amino acids. The number of identified phosphosites on serine (S), threonine (T), and tyrosine (Y) residues is shown using three different shades of blue.

(D) Violin plots showing the coefficient of variation (CV) of normalized phosphosite intensities across the analyzed groups. The black point indicates the median value of each distribution (median CV = 0.23, 0.24, and 0.22 for 2 h, 4 h, and 6 h, respectively).

(E) Principal component analysis (PCA) of the phosphoproteomics dataset showing the separation of samples across the analyzed conditions.

(F) Projection of the identified phosphosite clusters in PCA space, illustrating the relationships among the eight clusters.

(G) Kinase–substrate enrichment analysis predicting kinases responsible for phosphorylation of sites belonging to cluster 4 (NetworKIN).

(H) Mpk1 abundance across the analyzed time points.

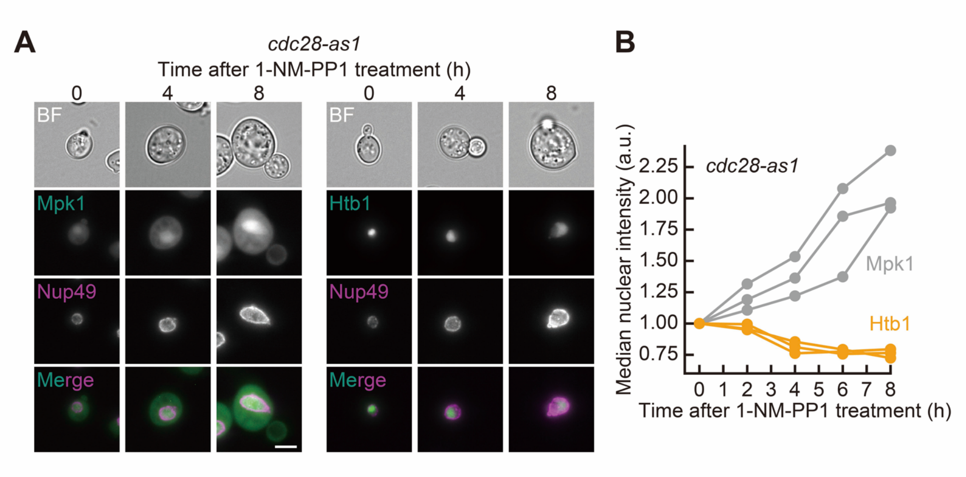


**Fig. S3. Observation of Mpk1 and Htb1 in *cdc28-as1* cells during excessive growth.**
(A) Representative images of Mpk1-mNeonGreen (left) and Htb1-mNeonGreen (right) in *cdc28-as1* cells after 1-NM-PP1 treatment, shown with bright field (BF) and Nup49-mScarlet-I images. Merged images of Mpk1- or Htb1-mNeonGreen with Nup49-mScarlet-I are also shown. For fluorescent protein signals, maximal intensity projection images are presented. Scale bar, 5 μm.
(B) Median nuclear Mpk1- or Htb1-mNeonGreen intensities in *cdc28-as1* cells following 1-NM-PP1 treatment. Values from three independent experiments are shown (n > 65 cells per time point per experiment). Values are normalized to the 0-h time point.

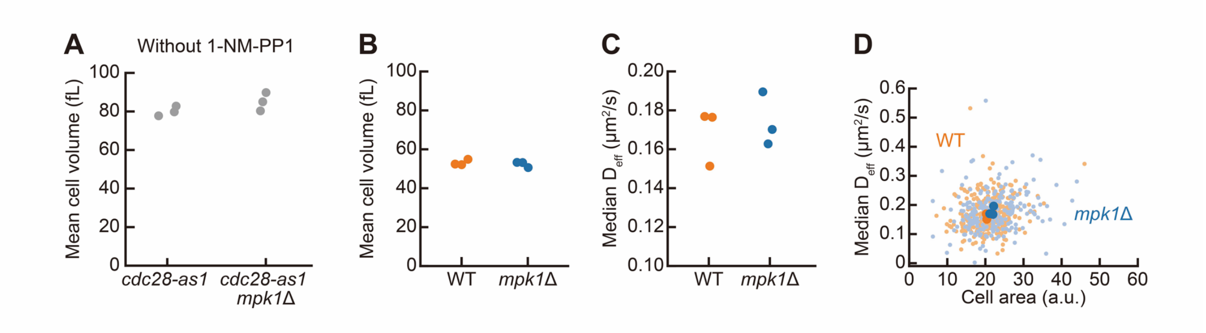


**Fig. S4. Particle mobility and cell volume in *cdc28-as1 mpk1*Δ, wild-type, and *mpk1*Δ strains.**

(A) Cell volume determined using a Coulter counter in *cdc28-as1* and *cdc28-as1 mpk1*Δ cells in the absence of 1-NM-PP1 from three independent experiments. Each dot represents the value from an individual experiment.

(B) Cell volume determined using a Coulter counter in wild-type (WT) and *mpk1*Δ cells from three independent experiments. Each dot represents the value from an individual experiment.

(C) Median $D_{eff}$ of 40nm-GEMs in WT and *mpk1*Δ cells from three independent experiments (n > 1700 trajectories for each experiment).

(D) Median $D_{eff}$ of 40nm-GEMs and cell areas at single-cell level in WT and *mpk1*Δ cells. Each small dot represents the median $D_{eff}$ of 40nm-GEMs in a single cell and the cell area measured in the same cell (n > 210 cells from three independent experiments). Each large dot indicates the median $D_{eff}$ of 40nm-GEMs and median cell area from each experiment. Cell areas were measured from bright field images.

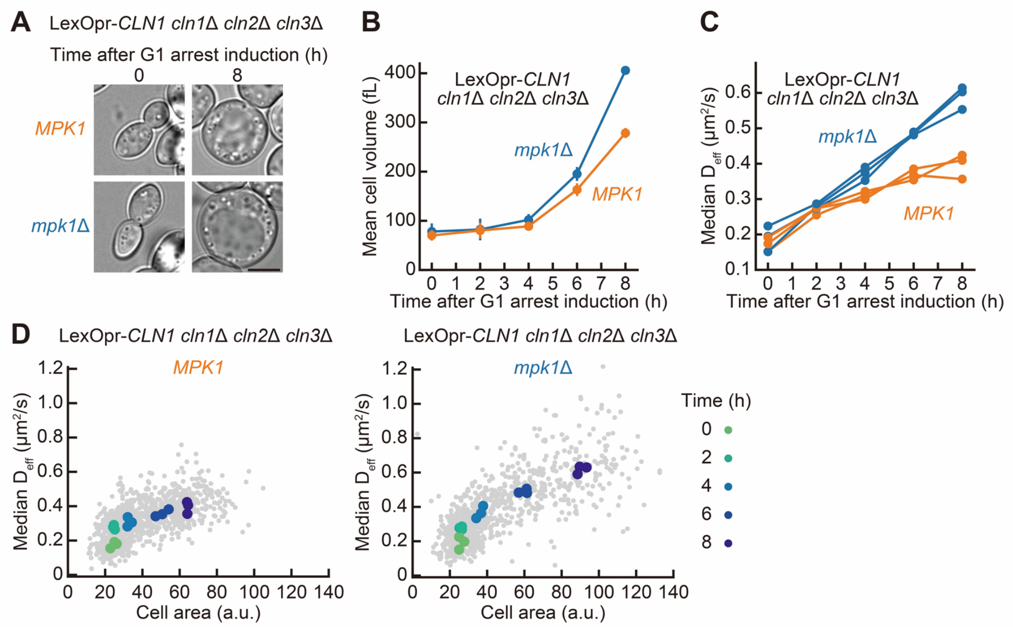


**Fig. S5. Particle mobility and cell volume in LexOpr-*CLN1 cln1*Δ *cln2*Δ *cln3*Δ strains.**

(A) Representative bright field images of LexOpr-*CLN1 cln1*Δ *cln2*Δ *cln3*Δ strains with (*mpk1*Δ) or without (*MPK1*) *MPK1* deletion after induction of G1 arrest. Scale bar, 5 μm.

(B) Cell volume determined using a Coulter counter in LexOpr-*CLN1 cln1*Δ *cln2*Δ *cln3*Δ strains after induction of G1 arrest. The mean values from the three independent experiments are represented with SD.

(C) Median $D_{eff}$ of 40nm-GEMs in LexOpr-*CLN1 cln1*Δ *cln2*Δ *cln3*Δ strains after induction of G1 arrest. The values from three independent experiments are indicated (n > 950 trajectories at each time point for each experiment).

(D) Median $D_{eff}$ of 40nm-GEMs and cell areas at single-cell level for LexOpr-*CLN1 cln1*Δ *cln2*Δ *cln3*Δ strains after induction of G1 arrest. Each small gray dot represents the median $D_{eff}$ of 40nm-GEMs in a single cell and the cell area measured in the same cell (n > 140 cells at each time point from three independent experiments). Each large dot indicates the median $D_{eff}$ of 40nm-GEMs and median cell area at each time point obtained from each experiment. Cell areas were measured from bright field images.

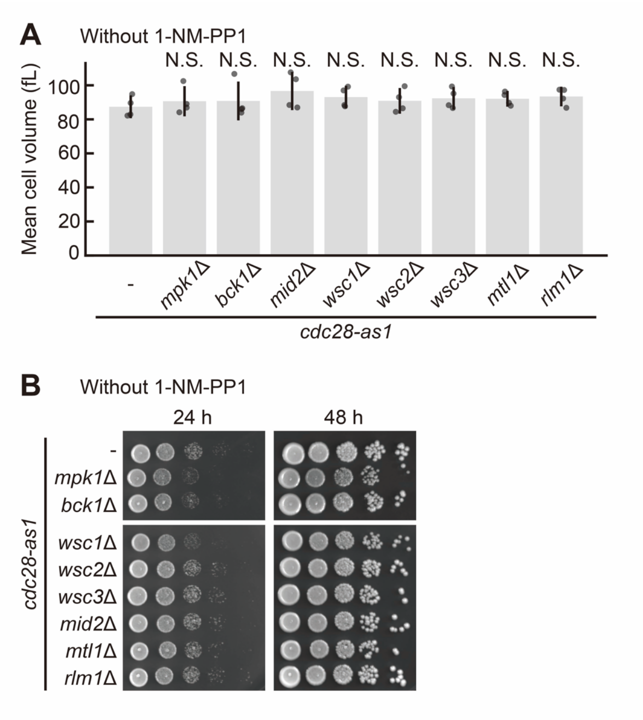


**Fig. S6. Characterization of double mutants of *cdc28-as1* lacking CWI pathway genes.**

(A) Cell volume determined using a Coulter counter in cells incubated without 1-NM-PP1. The mean values from the four independent experiments were represented with SD. Each dot represents the value from an individual experiment. Statistical significance was assessed by comparison with *cdc28-as1* (-) cells using one-way ANOVA followed by a *post hoc* Dunnett's test. N.S., not significant.

(B) Cells were cultured without 1-NM-PP1 and then 10-fold serial dilutions of the cells were spotted onto a YPD plate. The images were taken at 24 and 48 h after plating.

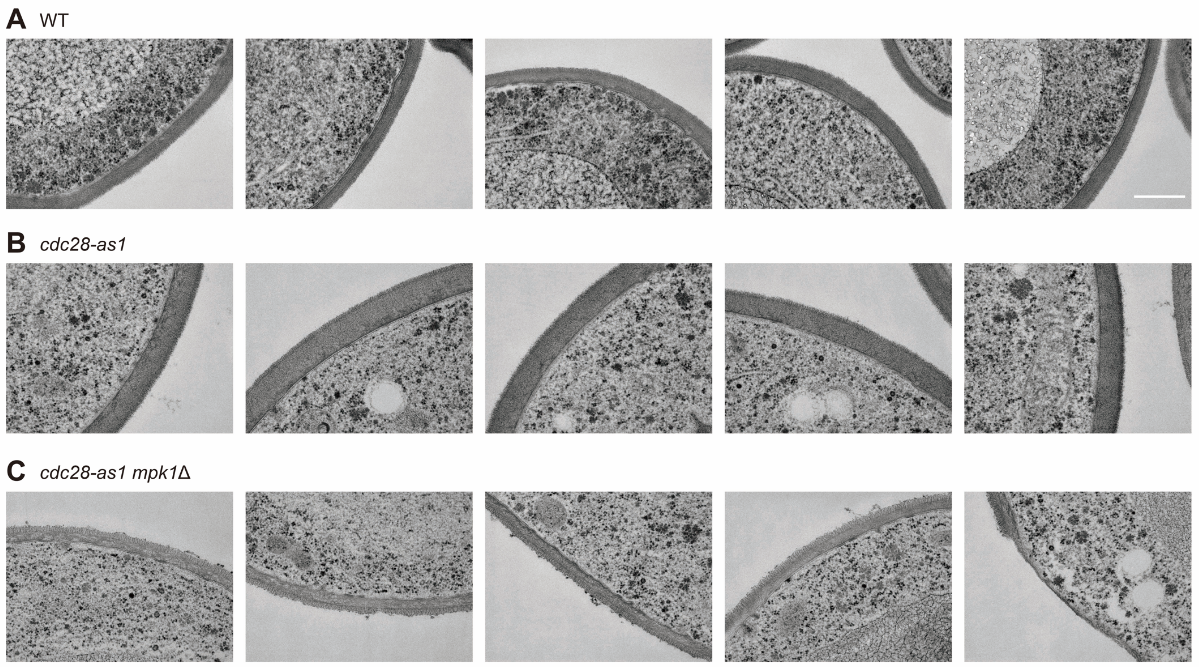


**Fig. S7. TEM images in wild-type, *cdc28-as1*, and *cdc28-as1 mpk1*Δ strains.**

(A-C) Representative TEM images in wild-type (WT) (A), *cdc28-as1* (B), and *cdc28-as1 mpk1*Δ (C) strains. Scale bar, 500 nm.

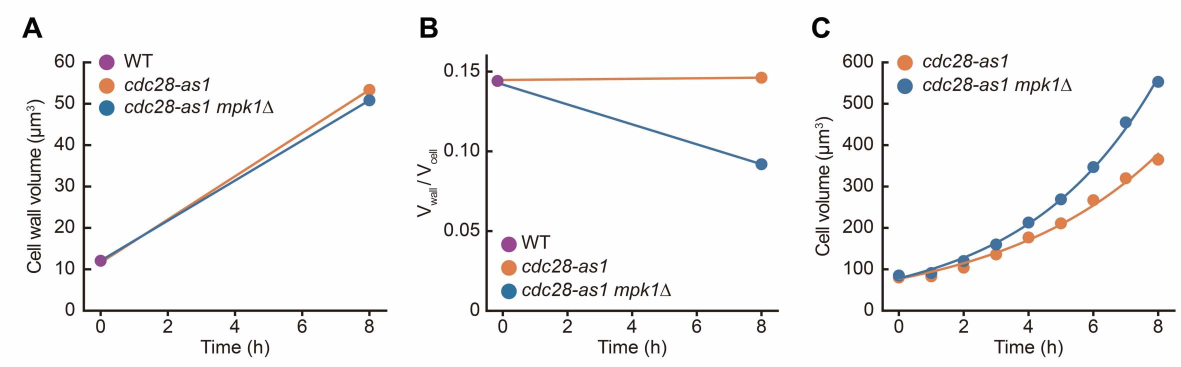


**Fig. S8. Cell wall volume, cell volume, and their ratio in *cdc28-as1* and *cdc28-as1 mpk1*Δ cells.**

(A) Cell wall volume increased at a similar rate over time for both *cdc28-as1* and *cdc28-as1 mpk1*Δ cells.

(B) The ratio of cell wall volume to total cell volume remained constant in *cdc28-as1* cells but decreased in *cdc28-as1 mpk1*Δ cells.

(C) Cell volume increased exponentially in both genotypes, with a higher expansion rate observed in *cdc28-as1 mpk1*Δ cells.

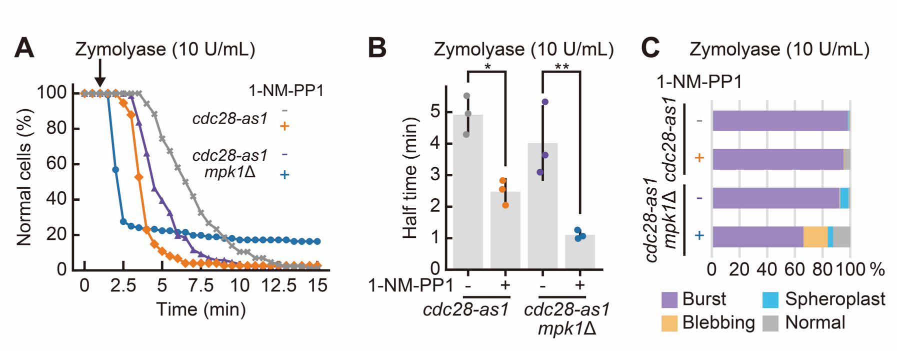


**Fig. S9. Sensitivity of *cdc28-as1* and *cdc28-as1 mpk1*Δ cells to Zymolyase (10 U/mL).**
(A) Proportion of normal cells after Zymolyase treatment (10 U/mL) during 15 min time course (n > 70 cells). *cdc28-as1* without 1-NM-PP1: gray, *cdc28-as1* with 1-NM-PP1: orange, *cdc28-as1 mpk1*Δ without 1-NM-PP1: purple, and *cdc28-as1 mpk1*Δ with 1-NM-PP1: blue.
(B) Half the time required for the proportion of normal cells to reach a plateau following Zymolyase treatment (10 U/mL). The mean values from the three independent experiments were represented with SD, and each dot represents the value from an individual experiment. Statistical significance was assessed using one-way ANOVA followed by a *post hoc* Tukey-Kramer test. *, $p < 0.05$; **, $p < 0.01$.
(C) Classification of cells after 14 min of Zymolyase treatment (10 U/mL). Cells were categorized into four types: burst, blebbing, spheroplast, and normal (n > 240 cells from three independent measurements).

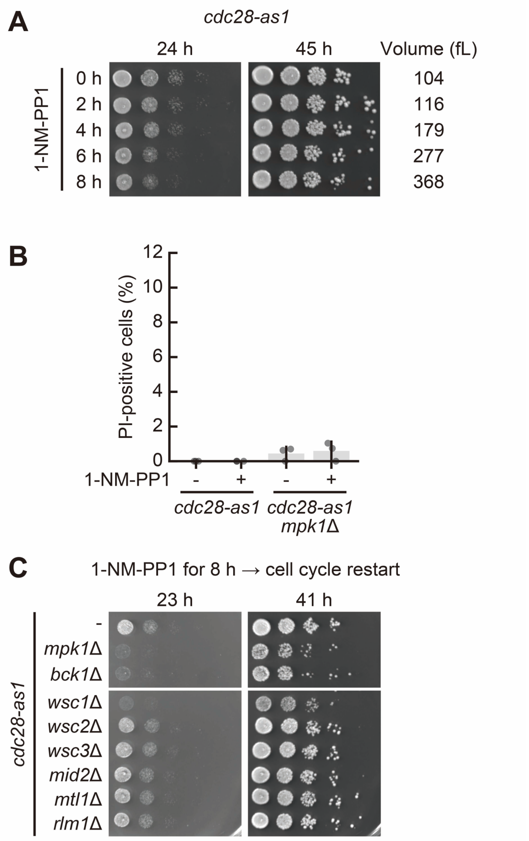


**Fig. S10. Characterization of cells before and after cell cycle restart.**

(A) *cdc28-as1* cells were incubated with 1-NM-PP1 for the indicated durations and then 10-fold serial dilutions of the cells were spotted onto a YPD plate. The images were taken at 24 and 45 h after plating. Mean cell volume at each time point is indicated.

(B) *cdc28-as1* and *cdc28-as1 mpk1*Δ cells were cultured with or without 1-NM-PP1 for 8 h and then stained with PI to detect dead cells. The mean values from the three independent experiments were represented with SD, and each dot represents the value from an individual experiment (n > 130 cells for each experiment).

(C) Cells were cultured in the presence of 1-NM-PP1 for 8 h and then 10-fold serial dilutions of the cells were spotted onto a YPD plate. The images were taken at 23 and 41 h after plating.

**Movie S1. The mobility of 40 nm particles in *cdc28-as1* cells during cell enlargement.**
Representative bright field (upper) and fluorescence (lower) images of *cdc28-as1* cells expressing 40nm-GEMs at the indicated time after 1-NM-PP1 treatment. The fluorescence images were taken every 20 milliseconds, and the movie playback speed is set to 50 frames per second. The timestamp shows time in seconds. Scale bar, 5 μm.

**Movie S2. The mobility of 40 nm particles in *cdc28-as1 mpk1*Δ cells during cell enlargement.**
Representative bright field (upper) and fluorescence (lower) images of *cdc28-as1 mpk1*Δ cells expressing 40nm-GEMs at the indicated time after 1-NM-PP1 treatment. The fluorescence images were taken every 20 milliseconds, and the movie playback speed is set to 50 frames per second. The timestamp shows time in seconds. Scale bar, 5 μm.

**Movie S3. Cellular responses of enlarged *cdc28-as1 mpk1*Δ cells to Zymolyase (10 U/mL).**
Representative time-lapse images of *cdc28-as1 mpk1*Δ cells treated with 1-NM-PP1 for 8 h followed by Zymolyase treatment (10 U/mL). Three types of cellular responses are shown: (1) burst, (2) blebbing, and (3) spheroplast. Zymolyase was added at 1 min. The movie playback speed is set to 5 frames per second. The timestamp is indicated as minutes and seconds. Scale bar, 5 μm.

**Movie S4. Cell cycle restart of enlarged *cdc28-as1* and *cdc28-as1 mpk1*Δ cells.**
Representative time-lapse images of *cdc28-as1* (left) and *cdc28-as1 mpk1*Δ (right) cells treated with 1-NM-PP1 for 8 h followed by the cell cycle restart. The movie playback speed is set to 5 frames per second. The timestamp is indicated as hours and minutes. Scale bar, 5 μm.
